# Chromatin run-on reveals the transcriptional etiology of glioblastoma multiforme

**DOI:** 10.1101/185991

**Authors:** Tinyi Chu, Edward J. Rice, Gregory T. Booth, H. Hans Salamanca, Zhong Wang, Leighton J. Core, Sharon L. Longo, Robert J. Corona, Lawrence S. Chin, John T. Lis, Hojoong Kwak, Charles G. Danko

**Affiliations:** Baker Institute for Animal Health, College of Veterinary Medicine, Cornell University, Ithaca, NY 14853.; Graduate field of Computational Biology, Cornell University, Ithaca, NY 14853.; Department of Biomedical Sciences, College of Veterinary Medicine, Cornell University, Ithaca, NY 14853.; Department of Molecular Biology and Genetics, Cornell University, Ithaca, NY 14853.; Department of Anesthesiology, SUNY Upstate Medical University, Syracuse, NY 13224.; Department of Molecular and Cell Biology, University of Connecticut, Storrs, Connecticut, USA; Department of Neurological Surgery, SUNY Upstate Medical University, Syracuse, NY 13224.; Department of Pathology, SUNY Upstate Medical University, Syracuse, NY 13224.

## Abstract

The human genome encodes a variety of poorly understood RNA species that remain challenging to identify using existing genomic tools. We developed chromatin run-on and sequencing (ChRO-seq) to map the location of RNA polymerase using virtually any input sample, including samples with degraded RNA that are intractable to conventional RNA-seq. We used ChRO-seq to develop the first maps of nascent transcription in primary human glioblastoma (GBM) brain tumors. Whereas enhancers discovered in primary GBMs resemble open chromatin in the normal human brain, rare enhancers activated in malignant tissue drive regulatory programs similar to the developing nervous system. We identified enhancers that regulate genes characteristic of each known GBM subtype, identified transcription factors that drive them, and discovered a core group of transcription factors that control the expression of genes associated with clinical outcomes. This study uncovers new insights into the molecular etiology of GBM and introduces ChRO-seq which can now be used to map regulatory programs contributing to a variety of complex diseases.

## Introduction

Our genomes encode a wealth of functional elements that play critical roles in the molecular basis of disease. RNAs serve as a marker for a surprisingly diverse group of functional elements, revealing the expression level of protein coding genes (mRNAs), as well as the location of enhancers and other non-coding regulatory elements which transcribe short and rapidly degraded non-coding RNAs (ncRNA)^1–5^. However, the discovery of ncRNA species, especially of enhancer-templated RNAs (eRNAs) characteristic of distal regulatory elements^2,5^, has proven challenging. Most ncRNAs are not represented in RNA-seq data, owing to the rapid degradation rates of most ncRNAs by the nuclear exosome complex^6,7^. Chromatin immunoprecipitation and sequencing (ChIP-seq) for RNA polymerase II is of limited value because it has a poor signal-to-noise ratio which obscures less abundant RNA species^8^. Likewise, assays that measure nuclease accessibility, such as DNase-I-seq^9^ and ATAC-seq^10^, are poor sources of information about transcriptional activity because they identify open chromatin regions irrespective of activity, and do not measure critical sources of information about mRNAs such as gene expression levels or transcript boundaries.

Recent studies have shown that sequencing nascent RNAs attached to an actively transcribing RNA polymerase complex is an effective strategy for discovering coding and ncRNAs^11–18^. Nascent RNA-seq techniques, such as Precision Run-On and Sequencing (PRO-seq)^13^, provide significantly higher sensitivity in detecting short-lived ncRNAs. Thus, PRO-seq and related assays provide a rich source of information about multiple layers of regulatory control, enabling simultaneous measurements of transcription at protein-coding genes and the discovery of active regulatory elements, including enhancers^19–21^.

Cancers are a particularly attractive target for nascent RNA sequencing techniques because cancer is a disease of gene regulation^22^. In most cancers, somatic changes to DNA sequence affect oncogenic or tumor suppressive pathways^23,24^. In some cases somatic mutations affect the core transcriptional machinery directly^25^, motivating the use of assays that directly measure the localization of Pol II. Somatic mutations initiate secondary changes in gene expression that are responsible for initiating changes in cell morphology and behavior that are characteristic of malignancy. For this reason, gene expression signatures from RNA-seq and other assays have proven effective as biomarkers, denoting cancer subtypes that are associated with progression and survival. However, which genes undergo regulatory changes in cancer, and especially the identity of key transcription factors that encode the malignant behaviors of cancer cells by their effect on target genes, remain poorly defined.

Nascent RNA sequencing techniques remain challenging to apply in some cell lines and especially to intact clinical isolates derived from cancer patients. Here we introduce a new chromatin-based run-on protocol, called Chromatin Run-On and Sequencing (ChRO-seq). ChRO-seq produces similar maps of transcription to PRO-seq in cell lines, but can also be applied to solid tissue samples, even those in which RNA is highly degraded. We used ChRO-seq to analyze 24 human glioblastoma multiforme (GBM) brain tumors, patient derived xenografts (PDXs), and a primary non-malignant brain sample. In addition to features of GBM already known from mRNA-seq data, ChRO-seq also revealed the location of thousands of promoters and distal enhancers that are active in primary GBM tissue. Analysis of rare distal enhancer elements suggests that primary tumors retain a surprising degree of similarity to the tissue of origin when grown *in vivo*. Nevertheless, we identified thousands of enhancers that change activity levels in tumors, providing new insights into the transcription factors responsible for malignant cell behavior. We also identified a core group of transcription factors that drive expression programs associated with poor clinical outcomes.

## Results

### Run-on assays in solid tissue

We developed Chromatin Run-On and sequencing (ChRO-seq), a new method to map RNA polymerase in cell or tissue samples (**Fig. 1a**). The primary challenge faced when using PRO-seq is often obtaining nuclei that are suitable for a run-on reaction. We therefore developed an alternative method which relies on fractionating insoluble chromatin, including engaged RNA polymerase II (Pol II)^27^ (see **Online Methods**). Insoluble chromatin was re-suspended by sonication and used as input to a run-on reaction (**Fig. 1a**). The run-on was designed to incorporate a biotinylated nucleotide triphosphate (NTP) substrate into the existing nascent RNA that provides a high-affinity tag used to enrich nascent transcripts. The biotin group prevents the RNA polymerase from elongating after being incorporated into the 3’ end of the nascent RNA when performed in the absence of normal NTPs, thus enabling up to single-nucleotide resolution for the polymerase active site^13,28^.

**Fig. 1.**
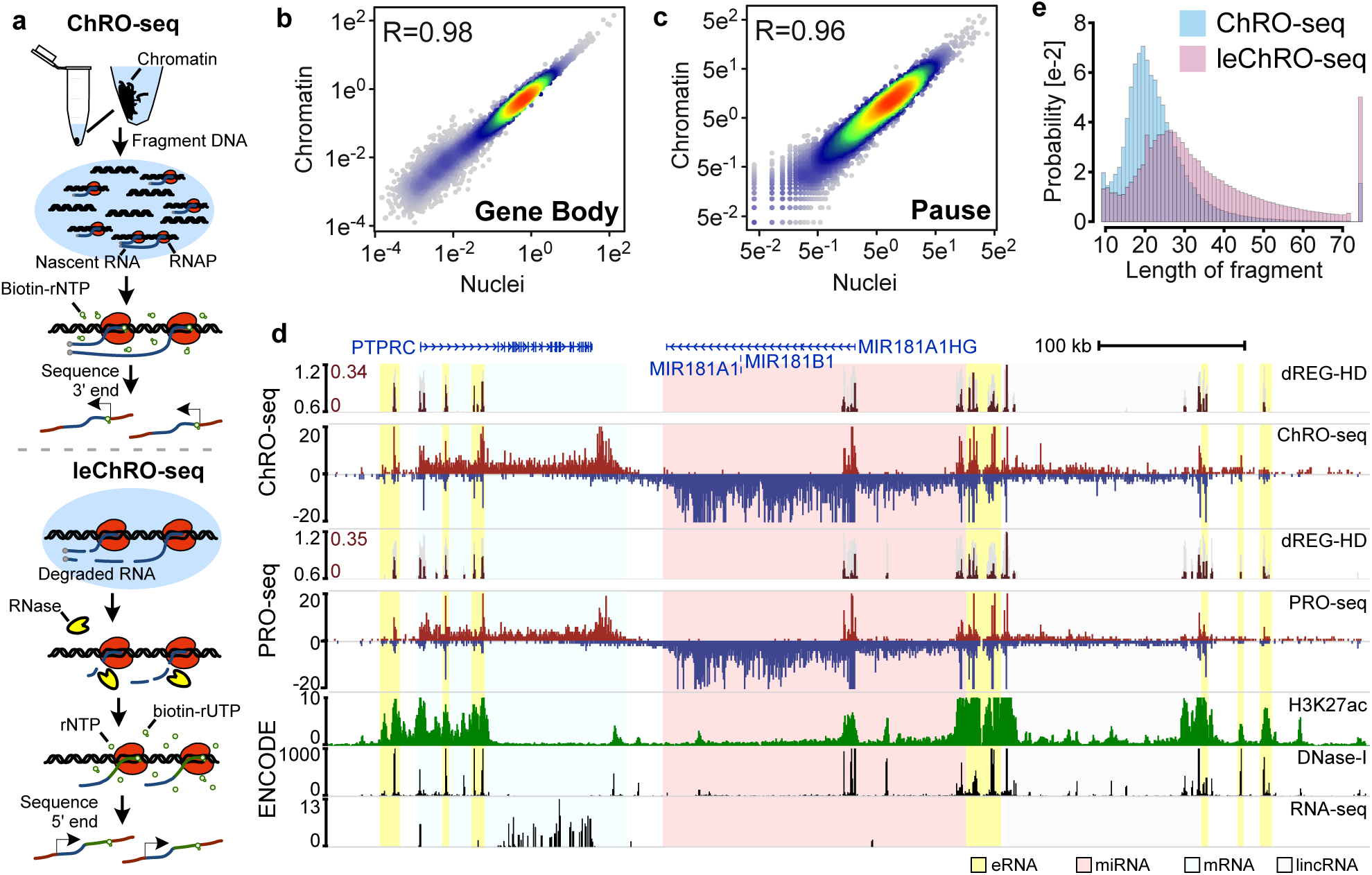
ChRO-seq and leChRO-seq measure primary transcription in isolated chromatin. (a) Isolated chromatin is incubated with biotinylated rNTPs, purified by streptavidin beads, and sequenced from the 3’ end. leChRO-seq degrades existing RNA, extends nascent transcripts an average of 100 bp, and sequences RNAs from the 5’ end. (b and c) Comparison between matched ChRO-seq and PRO-seq in annotated gene bodies (b) or at the peak of paused Pol II (c) in units of reads per kilobase per million mapped. (d) Comparison between ChRO-seq (top three tracks), PRO-seq (center), and H3K27ac ChlP-seq, DNase-I-seq, and RNA-seq (bottom). dREG-HD shows the raw signal fordREG (gray) and imputed DNase-I hypersensitivity signal (dark red). (e) The distribution of read lengths from ChRO-seq (blue) and leChRO-seq (pink) in a 30 year old primary GBM.

We performed matched ChRO-seq and PRO-seq experiments in the human Jurkat T-cell leukemia line, in which both nuclei and chromatin could be obtained. Median ChRO-seq signal across annotated genes was within the range of variation observed in PRO-seq data from the same cell line (**Supplementary Fig. 1**). In contrast, we noted differences in the pause peak and transcription past the polyadenylation site compared with mNET-seq and Nascent-seq, two other chromatin-based RNA sequencing assays^14,29,30^ (**Supplementary Note 1**). ChRO-seq and PRO-seq produced highly correlated levels of RNA polymerase in the bodies of mRNA encoding genes (R= 0.98; **Fig. 1b**). Likewise, signal for paused Pol II was highly correlated across the 5’ ends of annotated genes (R= 0.96; **Fig. 1c**), and pause levels in our test ChRO-seq library were within the range of variation observed using nuclei (**Supplementary Fig. 2**). The microRNA MIR181 locus illustrates the advantages of ChRO-seq compared with other molecular assays (**Fig. 1d**). Notably, both ChRO-seq and PRO-seq discovered the primary transcript encoding MIR181 as well as dozens of eRNAs that were not discovered using RNA-seq.

Because RNA prepared from archival tissues is often highly degraded, such samples are poor candidates for genome-wide transcriptome analysis using RNA-seq. The RNA polymerase-DNA complex is more stable than RNA^31^, suggesting that engaged polymerases may provide an avenue for producing new RNAs in archived samples. We obtained a primary glioblastoma multiforme (GBM) (grade IV, ID# GBM-88–04) that was stored in a tissue bank for 30 years. Bioanalyzer analysis confirmed that RNA was highly degraded in this sample (RIN = 1.0, **Supplementary Fig. 3**), thus precluding the application of RNA-seq (requires RIN of 2–4). To measure gene expression in this sample, we devised length extension ChRO-seq (leChRO-seq), a variant of ChRO-seq that uses transcriptionally-engaged Pol II and a mix of biotinylated-NTP and normal NTPs to extend degraded nascent RNA transcripts (**Fig. 1a**). Whereas libraries prepared without an extended run-on had a median insert size of 20 bp, precisely the length of RNA protected from degradation by the polymerase exit channel^32^, run-on samples achieved a longer RNA length distribution that was better suited for mapping unique reads within the human genome (**Fig. 1e**). Although RNA degradation could, in principal, destabilize RNA polymerase, we nevertheless observed that leChRO-seq produced maps of transcription that were correlated with those obtained using ChRO-seq and PRO-seq, suggesting that leChRO-seq accurately measures gene expression and pausing (**Supplementary Fig. 1a, 2, 4a**). Thus, leChRO-seq allows the robust interrogation of archival tissue samples which cannot be analyzed using standard genomic tools.

### Maps of transcription in primary GBMs

To demonstrate how ChRO-seq can provide insights into complex disease, we obtained ChRO-seq or leChRO-seq data from 20 primary glioblastomas, three patient derived xenografts (PDX), and a non-malignant brain (**Supplementary Table 1**). Histopathology revealed hallmarks of grade IV malignant astrocytoma in all GBMs (e.g., GBM-15–90, **Supplementary Fig. 5**). We sequenced ChRO-seq data from each GBM to an average depth of 33 million uniquely mapped reads per sample (10–150M reads/ sample). We confirmed that data collected from biopsies isolated from nearby regions (technical replicates) were highly correlated (**Supplementary Fig. 4c-f**, **Supplementary Note 2**). ChRO-seq data revealed changes in the transcription of several genes undergoing recurrent amplifications in GBMs^24,33^, including EGFR in GBM-15–90 (**Fig. 2a**).

**Fig. 2.**
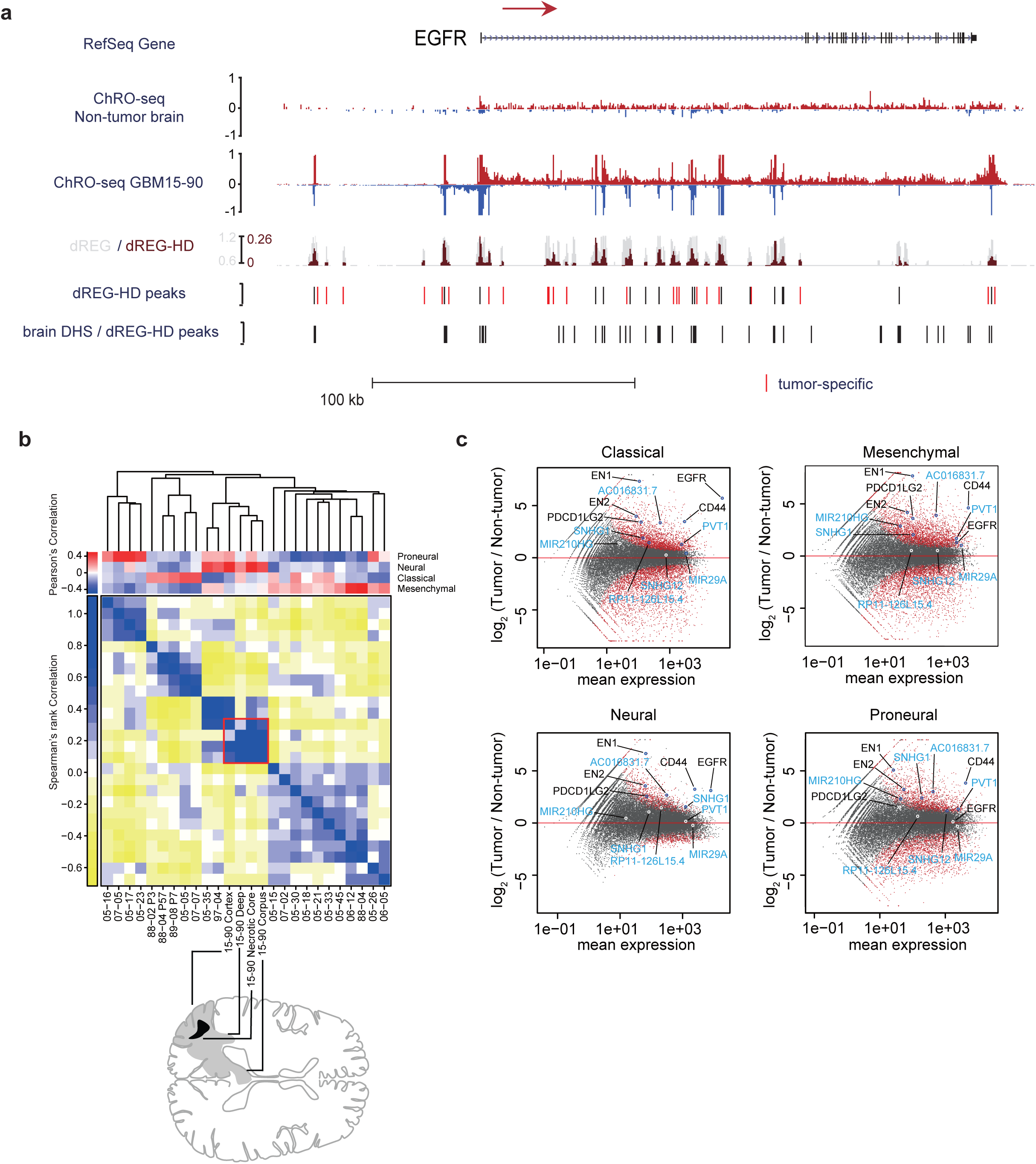
ChRO-seq detects transcription in primary human glioblastomas. **(a)** RPM normalized ChRO-seq signal at the EGFR locus in nonmalignant brain (top) and GBM1 (center). dREG (gray) and dREG-HD (dark red) signals are shown for GBM-15–90 (track 3). dREG-HD sites that are not DHSs in adult brain reference samples are highlighted in red (track 4). DHSs in 6 adult brain reference samples and dREG-HD peaks from the nonmalignant brain sample (track 5). **(b)** Upper matrix: subtype scores for each patient, calculated by Pearson’s correlation with the centroid of gene expression of corresponding subtype. Lower matrix: Spearman’s rank correlation in 20 primary GBMs representing 840 signature genes. Red square denotes four regions dissected from GBM-15–90. Sample order is based on single-link hierarchical clustering of the lower matrix, shown by the dendrogram. **(c)** Differential gene transcription of primary GBMs in each subtype compared with non-malignant brain. Genes of interest are highlighted. lncRNAs are highlighted in blue.

To gain further insight into how transcription changes in malignant tissue, we analyzed transcription within annotated protein-coding genes and non-coding RNAs. GBMs from our cohort represent each of the four previously reported molecular subtypes^26^ (**Fig. 2b**, **Supplementary Fig. 6**). Though most tumors have transcription patterns characteristic of one dominant molecular subtype, several tumors in our cohort were similar to multiple subtypes, especially those matching neural and mesenchymal signatures, consistent with reports of cellular heterogeneity within the same tumor^34,35^ (**Fig. 2b**). We identified 3,504 protein-coding genes and 1,250 ncRNAs that were differentially transcribed across all 20 primary GBMs relative to replicates of the non-malignant brain (*p* < 0.05, False discovery rate [FDR] corrected, DESeq2^36^). Differentially transcribed genes had notable enrichments in biological processes related to cell cycle, DNA replication / metabolic processes, development (up-regulated in the tumor), and nervous system homeostasis (down-regulated) (**Supplementary Fig. 7**). For example, multiple transcription factors with a role specifying nervous system development were expressed more highly in nearly all tumors, including the *HOX* gene clusters and engrailed-1 and 2 *(EN1* and *EN2)* (**Fig. 2c**; **Supplementary Fig. 8**). Notably, we discovered several differentially transcribed long non-coding RNAs (lncRNAs) that confer growth advantages to U87 glioblastoma cells^37–40^ (e.g., AC016831.7, PVT1, SNHG1, etc. **Fig. 2c**; **Supplementary Table 2**). Taken together, our analysis of ChRO-seq data identified transcriptional changes common among all GBMs in our cohort, many of which were consistent with previous analyses of primary GBMs based on the abundance of mRNA, as well as differentially transcribed lincRNAs that may have clinical value.

### GBM enhancers retain signatures of normal brain tissue

Active transcriptional regulatory elements (TREs), including promoters and enhancers, have a characteristic pattern of RNA polymerase initiation that allows their discovery using ChRO-seq data^2,5,17,19–21^. We developed a novel algorithm to identify the precise location of active TREs, called dREG-HD, which takes PRO-seq or ChRO-seq data as input and identifies TREs that are similar to the subset of DNase-I hypersensitive sites (DHSs) that exhibit local transcription initiation. The dREG-HD algorithm improved the resolution of dREG^19^ by imputing smoothed DNase-I-seq signal intensity, and identified sites initiating transcriptional activity with 80% sensitivity at >90% specificity (**Supplementary Fig. 9**). dREG-HD recovered the nucleosome depleted region in histone modification ChIP-seq and MNase-seq data (**Supplementary Fig. 10**), demonstrating that it had substantially higher resolution compared with dREG alone.

The vast majority (96%) of TREs identified by dREG-HD in each primary GBM sample were DHSs in at least one of the 216 reference tissues analyzed by ENCODE or Epigenome Roadmap^41,42^. However, most DHSs were discovered in only a few of the tissues in the reference dataset (**Fig. 3a**) and were distal (>1 kb) to annotated transcription start sites (**Fig. 3b**), suggesting that many reflect the activity of cell-type specific distal enhancers in the tumor. Rare distal TREs (henceforth referred to as "enhancers”) provide a unique “fingerprint” for quantitatively evaluating the similarity between two samples, and could be used to define the relationship between tumors and normal tissue.

**Fig. 3.**
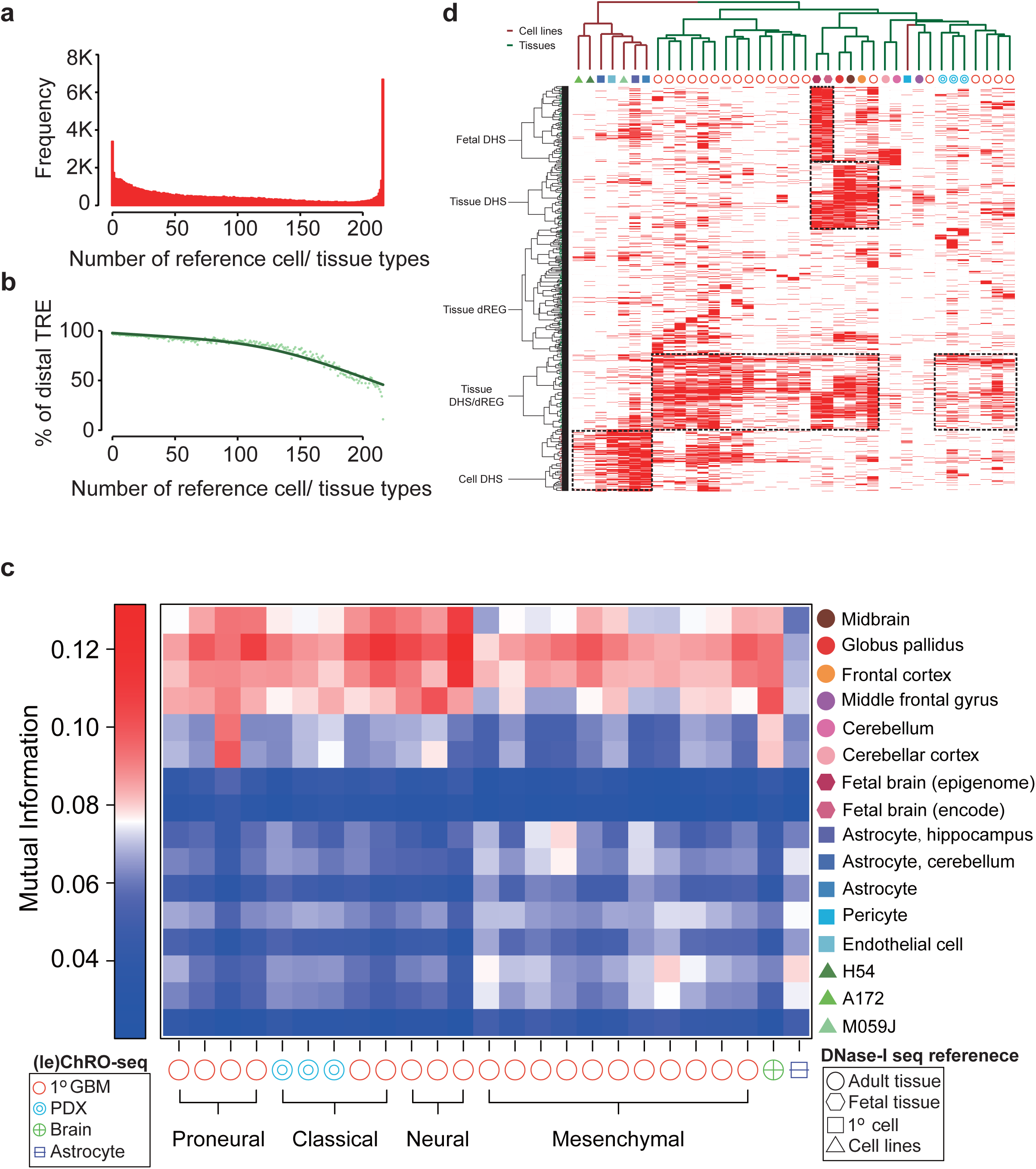
Comparison between TREs in primary GBM / PDX and reference DHSs. **(a)** Histogram representing the number of reference samples that have a DHS overlapping each dREG-HD site found in any of the 23 primary GBM / PDX samples. **(b)** Percentage of TREs >1kb from the nearest GEN-CODE transcription start site. **(c)** Mutual information between TREs in the indicated GBM and refer-ence sample. **(d)** Clustering of reference samples with primary GBM / PDX based on the activation of TRE. Activate TREs are marked in red; inactive ones are in white.

We developed a strategy that compares active enhancer landscapes obtained using dREG-HD with DHSs across all public datasets (see **Online Methods**). Our strategy consistently discovered the expected cell lines (**Supplementary Fig. 11**), even identifying the expected genotype (GM12878) among all lymphoblastoid cell lines as the most similar to GM12878 PRO-seq data (**Supplementary Fig. 11b**). Using unique enhancers to “fingerprint” primary GBM samples revealed enhancer landscapes that were highly similar to normal brain reference samples compared to other reference tissues (**Fig. 3c**, **Supplementary Fig. 12**). In GBM-15–90, for instance, 86% of TREs were shared with primary brain tissue, which was greater similarity than observed in either GBM cell lines (62% TRE identity) or *in vitro* cultured primary brain cells (75%) (**Supplementary Fig. 13**).

To evaluate whether contamination of the GBM with normal brain tissue explained the extensive similarity with normal brain reference samples, we used leChRO-seq data from three PDXs, in which primary GBMs were grown in a murine host. In PDXs, murine cells replace both normal tissue and stroma^43^, and can be distinguished from tumor cells based on species-specific differences in DNA sequence. Mutual information ranked all PDX samples as similar to the normal human brain compared with all other samples (**Fig. 3c**). Thus we conclude that primary GBM cells are more similar to their cell of origin than may have been anticipated based on prior analysis of cell models.

Two models might explain differences in enhancer profiles between primary and cultured GBM cells. Differences might reflect either evolutionary changes as cancer cells adapt to *in vitro* tissue culture conditions, or differences in the cellular microenvironment between tissue culture and primary tumors. To distinguish between these two models, we used TREs to cluster 20 primary GBMs, 3 PDXs, 8 normal brain tissues, 3 GBM cell lines, and 5 brain-related primary cell types which were dissociated from the brain and grown *in vitro* for a limited number of passages. This analysis supported two major clusters, one composed of normal brain and tumor tissues grown *in vivo* and the other of cells grown *in vitro* (**Fig. 3d**, **Supplementary Fig. 14**). Notably, PDX samples clustered with the primary brain samples, demonstrating that PDXs are a reasonably accurate model for many of the transcriptional features associated with primary tumors. That primary brain cells passaged for a limited duration in tissue culture clustered with the GBM models strongly implicates the microenvironment in causing differences in the enhancer landscape of cells.

### TREs define three distinct regulatory programs activated in GBM tissue

TREs that were active in tumor tissue, but were not DHSs in any of the available adult brain reference samples, are strong candidates for contributing to the malignant phenotype of the tumor. Such tumor-associated TREs (taTREs) comprised 2–24% of TREs in each tumor (**Supplementary Fig. 15, 16, Supplementary Table 3**). We developed a statistical test to identify tissues which shared unexpectedly high overlap with taTREs identified in each tumor that controls for DHS scarcity (**Supplementary Table 4**) (see **Online Methods**). Hierarchical clustering of the taTREs among significant cell types revealed three regulatory programs that were enriched in the primary GBMs; one resembling a stem-like regulatory program, one associated with differentiated support cells, and a cluster of immune cells (**Fig. 4a**, **Supplementary Fig. 17**). taTREs significantly (*p* < 1e-4, bootstrap test) overlapped DHSs in fetal tissues of the nervous system (2.3–6.6-fold enrichment in 11/ 23 GBMs), especially spinal cord and brain, two fetal tissues derived from the neuroectoderm (**Fig. 4a**, see "Outlier tissues”). We also found evidence for enrichment in additional developmental tissues, for example embryonic stem cells and other fetal tissues from a variety of germ layers, and for a number of terminally differentiated support cell lineages including astrocytes, endothelial cells, fibroblasts, and osteoblasts. We emphasize that activation of these separate transcriptional regulatory programs may reflect gene expression changes in subsets of cells within the tumor. Overlap between taTREs and fetal brain tissue likely reflects the activation of a regulatory program that promotes stem-like properties observed in a population of GBM cells^44^. Similarly, overlap with astrocytes, endothelial cells, fibroblasts, or osteoblasts may capture tumor cells that have trans-differentiated into these lineages^45,46^. Notably, these two signatures were detected in PDX samples as well as primary GBMs, demonstrating that these signatures reflect transcriptional diversity in malignant cells.

**Fig. 4.**
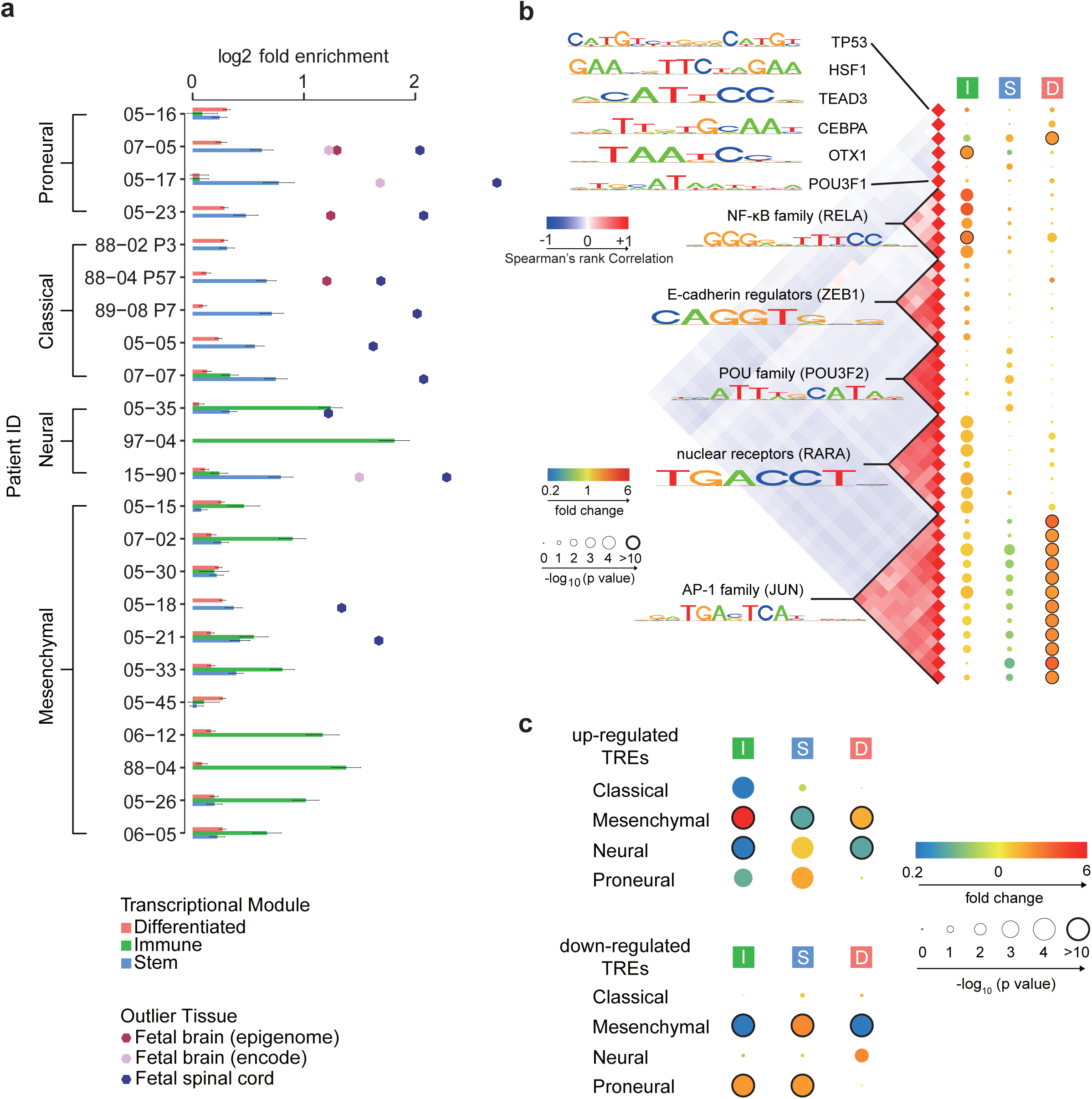
Tumor associated TREs (taTREs) activate three regulatory programs. **(a)** Barplots show the fold enrichment of reference tissues in the corresponding GBM. Reference samples were grouped into three clusters, representing stem-like (blue), immune (green), and differentiated (pink) regulatory programs. Error bars represent the standard error. Outliers with 6 times the standard error are highlighted. **(b)** Transcription factor binding motifs enriched in TREs that are members of the indicated regulatory program compared with TREs active in the normal brain. All motifs shown were significantly enriched following Bonferroni adjustment of the threshold p-value in at least one patient (*p* < 0.05 / 1882). The correlation heatmap (left) shows the correlation in DNA sequences recognized by motifs. Blue denotes a negative correlation and red denotes a positive correlation. Families of transcription factors and their representative motifs are highlighted. The median p value across patients significantly enriched/depleted (*p* < 0.05) in taTREs for each motif (right) are represented by the radius of the circle and enrichment (red) or depletion (blue) are represented by the color. **(c)** The enrichment of the indicated regulatory programs in subtype-biased TREs. The p value of enrichment/depletion for each regulatory program is represented by the radius of the circle and enrichment (red) or depletion (blue) are represented by the color.

To identify transcription factors involved in maintaining each regulatory program, we classified the taTREs in each tumor sample into regulatory programs based on their cell type overlap, and searched for enriched transcription factor binding motifs (*p* < 0.05 / 1882 in at least one patient, Fisher’s exact test, Rtfbsdb^47^). As we were limited in our ability to distinguish between paralogous transcription factors that share similar DNA binding specificities, we clustered motifs into 11 distinct groups, each associated with multiple transcription factors that may contribute to differences in expression (**Fig. 4b**). Many of these motifs showed mutually exclusive enrichment in the three regulatory programs (**Fig. 4b**; **Supplementary Fig. 18**), supporting the hypothesis that each regulatory program is a transcriptionally distinct program mediated by a different group of transcription factors. We identified POU domain containing transcription factors enriched in taTREs in the stem-like regulatory program. To verify that this enrichment reflects bona-fide binding of the predicted transcription factor, we obtained ChIP-seq data from cultured glioma neurospheres^44^. As predicted, taTREs in the stem-like program were enriched in both ChIP-seq reads and peak calls for POU3F2 (**Supplementary Fig. 19,20**). The differentiated support cell program was highly enriched for binding of activating protein 1 (AP-1), a heterodimer of the transcription factors FOS and JUN, a motif resembling heat shock factor 1 (HSF1), and the TEAD family (**Fig. 4b**). The immune program was enriched for C/EBP family (C/EBPA), NF-κB family (RELA), and the Retinoic Acid Receptor family (RARA), in agreement with reports that at least two of these factors play an important role in inflammatory responses in GBM^48,49^. Taken together, we have identified taTREs that correlate with complex behaviors intrinsic to malignant cells, for instance the stem-like regulatory program that was shared with neuroectodermal tissue, and identified candidate transcription factors that contribute to each behavior.

We asked how the stem, immune, and differentiated regulatory programs relate to previously described molecular subtypes in GBM. We used ChRO-seq signal to identify 6,775 TREs that were differentially transcribed in 2–3 primary GBMs most characteristic of each molecular subtype relative to samples representing the other three subtypes (*p* < 0.01, DESeq2; **Supplementary Table 4**). We compared subtype-biased TREs with those in the stem, immune, and differentiated regulatory program. TREs upregulated in mesenchymal GBMs were enriched 6-fold in the immune regulatory program (*p* < 1e-10, Fisher’s exact test; **Fig. 4c**), consistent with the mesenchymal subtype having higher numbers of tumor infiltrating immune cells^35,48^. TREs up-regulated in neural and proneural GBMs were enriched in signatures in the stem-like program (**Fig. 4c**). Nevertheless, TREs in the stem, immune, and differentiated regulatory programs did not always correlate with molecular subtype. For instance, two of the neural tumors in our cohort had a substantial immune regulatory program, and several mesenchymal tumors were strongly enriched for a stem-like program (**Fig. 4a**). Thus, the three regulatory programs discovered on the basis of rare enhancer fingerprints were distinct from previously reported subtypes, motivating correlations between these clusters and clinical outcomes once larger cohorts of tumors are analyzed using ChRO-seq.

### Transcription factors controlling GBM subtype

Transcriptional heterogeneity among GBMs is established in large part by the differential activity of transcription factors. To identify transcription factors that are involved, we focused on TREs with evidence of expression changes among the four previously described molecular subtypes (*p* < 0.01, DESeq2). We identified 38 binding motif clusters with extremely strong evidence of enrichment in active TREs with biased transcription in any subtype (*p* < 0.05 / 1882, Fisher’s exact test, **Fig. 5a**). Significantly enriched motifs passing our stringent multiple testing correction threshold were most common in the mesenchymal and neural subtypes, in which several had previous support in the literature, including those recognized by nuclear factor-κB (NF-κB) family and CCAAT/Enhancer Binding Protein (C/EBP) family enriched in TREs up-regulated in mesenchymal tumors^48,49^. Additionally, we identified numerous novel motif associations that correlate with subtype-biased expression including, for instance, RARA, SRF, SOX-family, and FOX-family.

**Fig. 5.**
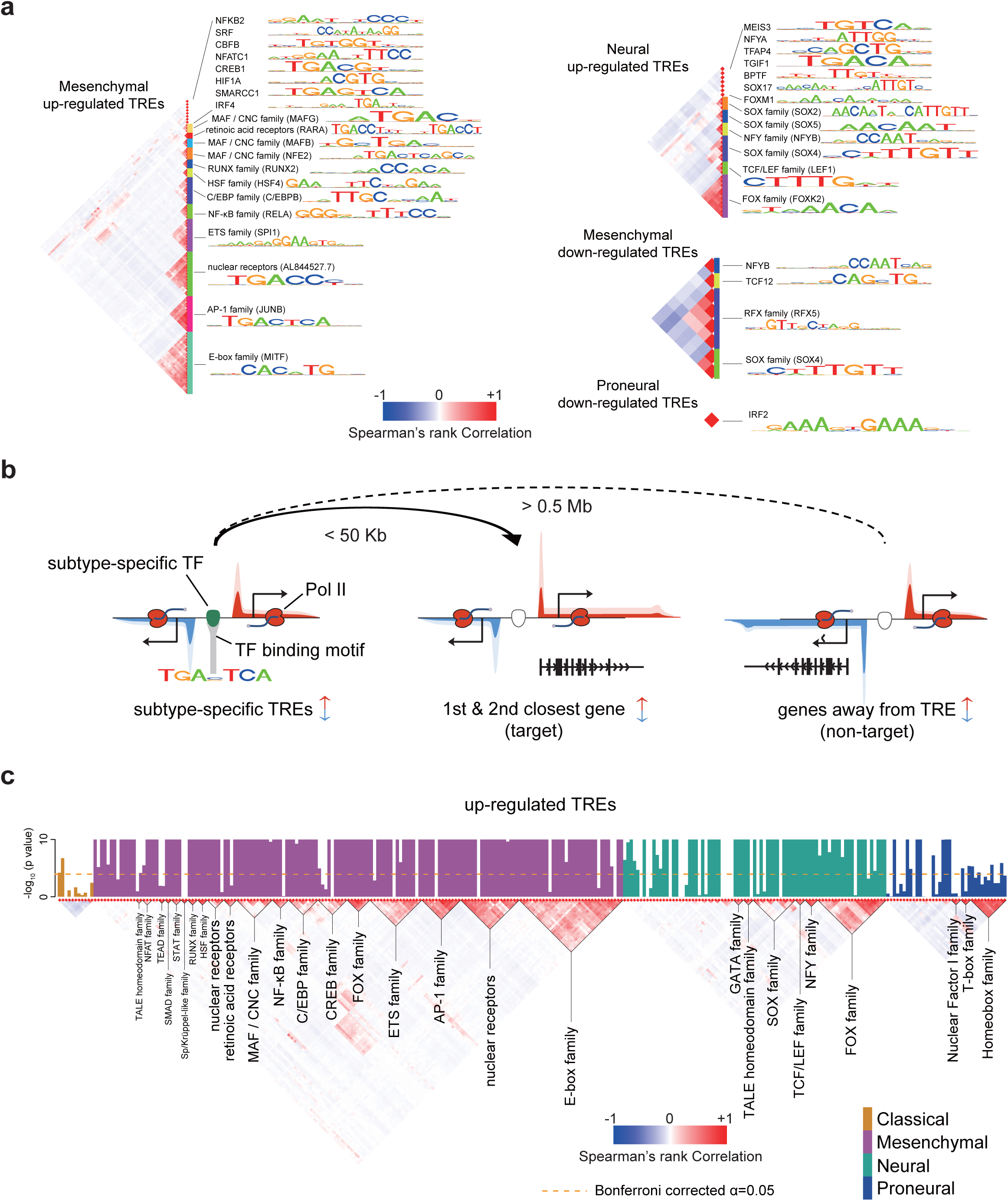
Transcription factors influencing transcriptional heterogeneity in GBM. **(a)** Transcription factor binding motifs enriched in TREs that were up- or down-regulated in the indicated subtype. All motifs shown were significantly enriched following Bonferroni adjustment of the threshold p value (*p* < 0.05 / 1882). The correlation heatmap (left) shows the correlation in DNA sequences recognized by motifs. Families of transcription factors and their representative motifs are highlighted. **(b)** Cartoon depicting the heuristics used to identify target genes of subtype-specific transcription factor and for defining non-target (control) genes. Changes in transcription of both target and non-target genes are of the same direction as that of subtype-biased TREs. Target genes are the 1st and 2nd genes within 50 Kb of the TRE. Non-target genes are at least 0.5 Mb away. **(c)** Barplots show the -log10 Wilcoxon rank sum p value of having higher correlations among target genes of each transcription factor binding motif (columns). Barplots are colored by subtype in which they were found to be enriched (p < 0.05, Fisher’s exact test). The correlation heatmap (bottom) shows the correlation in DNA sequence recognized by each motif. Transcription factor families are indicated below the plot. The dotted line shows the Bonferroni adjusted threshold for the between-target validation experiment.

Next we set out to identify target genes regulated by each transcription factor in GBM cells. First, we assume that molecular subtypes described in current literature do not completely describe the full range of heterogeneity among GBMs. To identify motifs contributing to heterogeneity that are only weakly correlated with the known molecular subtypes, we relaxed our statistical cutoff to a more permissive threshold at which we expected substantially higher sensitivity at an acceptable false discovery rate (*p* < 0.05, nominal Fisher’s exact test, **Supplementary Fig. 21, see Online Methods**). We identified bound occurrences of each enriched motif using heuristics that provide substantial performance improvements over existing high-resolution tools^50^. Motif occurrences were connected with the closest two annotated genes sharing similar subtype-bias within 50 kb (**Fig. 5b**), using fairly stringent heuristics to limit false discovery rates (see **Online Methods**). As expected, changes in transcription of TREs correlated with nearby genes, and were strongest for the nearest 1–2 genes from each TRE (**Supplementary Fig. 22**). Moreover these changes in the nearest two genes explained many of the markers defined in microarray studies^26^ (**Supplementary Fig. 23**).

To validate motifs and predicted target genes, we used the expectation that genes which share a common transcription factor should have expression levels that are more highly correlated with one another across tumors. We analyzed an independent RNA-seq dataset from a cohort of 174 primary GBMs^24^. Among the 304 transcription factors enriched in any subtype we noted a significantly stronger correlation between putative target genes for 235 (77%) compared with randomly selected genes matched for similar subtype specificity (**Fig. 5c**; **Supplementary Fig. 24a**). Furthermore, in two cases (NF-κB and STAT1), we found PRO-seq or RNA-seq data following activation of a signaling pathway targeting that transcription factor^51,52^. Despite both published experiments occurring in a different cell type and environmental context, we nevertheless found predicted targets to be 3.0-fold (NF-κB; *p* < 3.0e-21, Fisher’s exact test) and 6.9-fold (STAT1, *p* = 1.9e-11, Fisher’s exact test) enriched in genes responding in these experiments. Thus we have identified transcription factors contributing to major GBM expression subtypes, and a set of putative target genes of each transcription factor.

### Direct inference of transcription factor regulatory activities in GBMs

The gene-regulatory “trans” activities that a transcription factor has on its complement of bound TREs can be regulated by multiple transcriptional and post-transcriptional mechanisms. While some transcription factors are controlled predominantly by the abundance of its protein, many require a subsequent step such as post-transcriptional activation of the protein product to regulate target genes (**Fig. 6a**). We asked whether we could distinguish between these two broad regulatory activities by using ChRO-seq, and using an integrative analysis incorporating both ChRO-seq and publicly available mRNA-seq data.

**Fig. 6.**
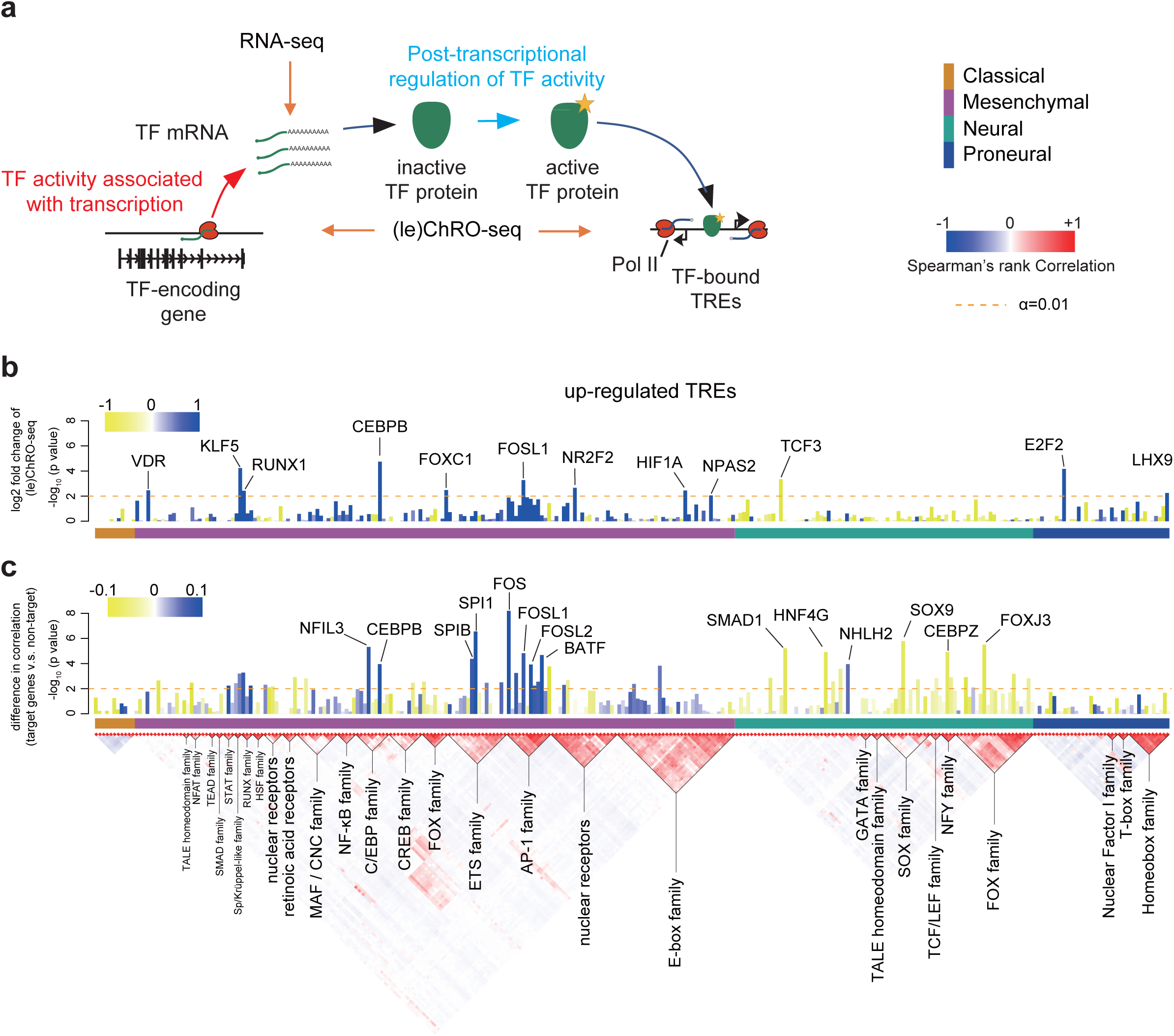
Regulatory activities of transcription factors are controlled by transcription and post-transcriptional mechanisms in GBM. **(a)** The cartoon illustrates the various stages at which transcription factor activities can be regulated and the corresponding signals detected by RNA-seq and (le)ChRO-seq. The activity of some transcription factors correlates predominantly with the abundance of its protein. Many transcription factors require post-transcriptional activation of the protein product before regulating target genes. **(b)** Barplot shows the FDR corrected -log10 p value (DESeq2) representing changes in Pol II abundance detected by (le)ChRO-seq on the gene encoding the indicated transcription factor. The level of upregulation (blue) and downregulation (yellow) is indicated by the color scale (log-2). The horizontal color bar below the barplot indicates the corresponding subtype in which the motif shows enrichment in the upregulated TREs. The dashed line shows the the FDR corrected α at 0.01. **(c)** The barplot shows the -log10 Wilcoxon rank sum test p value denoting differences in the distribution of correlations between the mRNA encoding the indicated transcription factor and either target or non-target control genes. The difference between the median correlation of target and non-target genes is indicated by color. Blue represents higher correlation between mRNA and target genes and yellow indicates a lower correlation. Dashed line shows the the uncorrected α at 0.01. Outliers are labeled.

In the simplest mode of regulation, the gene-regulatory activity of a transcription factor is determined by the abundance of its protein, which can be correlated with the transcriptional activity of its gene and the abundance of its mRNA. To detect this type of regulatory activity using ChRO-seq data, we asked whether motifs enriched in active TREs of each subtype correspond to changes in Pol II density on the primary transcription unit encoding any one of the transcription factors that recognize the corresponding binding motif. In some cases, we observed transcriptional changes in the transcription factor coding gene in the same subtype in which we also observed motif enrichment (**Fig. 6b**; **Supplementary Fig. 24b**). Likewise, we found several cases in which mRNA encoding each transcription factor was correlated with the expression of its putative target genes across GBMs to a greater extent than expected based on a null model that controls for molecular subtype (**Fig. 6c**; see **Online Methods**). When we observed correlated changes, Pol II (or mRNA abundance) on the transcription factor coding gene typically changed in the direction expected given the known activating or repressive properties of that transcription factor. For instance, ChRO-seq signal in the gene body encoding the transcriptional activator CEBPB increased by 4.88-fold in mesenchymal tumors (**Fig. 6b**), consistent with a 2.43-fold enrichment of its corresponding motif in mesenchymal upregulated TREs (**Fig. 5a**).

We devised a strategy to estimate which transcription factors have gene-regulatory activities that were regulated by transcriptional or post-translational mechanisms. Focusing on the 25 unique motifs enriched in up-regulated TREs that are associated with multiple transcription factors, we found evidence of correlated changes in ChRO-seq data for eight (**Fig. 6b**). Likewise, 16 transcription factor families had a significantly higher correlation between the transcription factor mRNA and its putative target genes across available RNA-seq datasets than expected by a null model controlling for molecular subtype (**Fig. 6c**). Several of these correlations were weak in magnitude, which may be consistent with gene-regulatory activities controlled by multiple regulatory mechanisms for these transcription factors. We conservatively identified at least six transcription factors, including TEAD, GATA, HSF, NF-kB, and other transcription factor families, which had low correlations with their putative targets in RNA-seq and no evidence of transcriptional changes in ChRO-seq. These transcription factors were regulated primarily at a post-transcriptional level in GBM. For these transcription factors, ChRO-seq is an especially rich source of information about gene-regulatory activities.

### Transcription factors control groups of survival-associated genes in mesenchymal GBMs

Known molecular subtypes of GBM do not correlate with survival^26^, presenting a motivation to identify new classifiers that may have prognostic value. We hypothesized that the activity of transcription factors which control transcriptional heterogeneity among GBM patients may control biological functions correlated with survival. To determine whether gene-regulatory activities of transcription factors may be useful in predicting clinical outcomes, we compared the hazards ratio at putative target genes of each subtype specific binding motif. We analyzed two sets of non-target control genes: 1) The nearest annotated transcription start site (within 50 kb) of each subtype-specific TRE that was not changed in that subtype, and 2) Differentially transcribed genes in the same subtype that were not identified as targets, because the transcription start site was >0.5Mb away from the nearest putative binding site. Our analysis identified six transcription factors significantly associated with poor clinical outcomes, all in mesenchymal tumors (*p* < 0.05 / 432, Wilcoxon, **Fig. 7a**, **Supplementary Fig. 25**), which we clustered into three unique DNA binding specificities (RAR, C/EBP family, and RELA [NF-κB] **Supplementary Fig. 26**). Only one of these transcription factors, C/EBP, was associated with survival at the mRNA level (**Supplementary Fig. 27**), consistent with the gene-regulatory activity of C/EBP family correlating with the abundance of its mRNA (**Fig. 6b**). RELA activity was correlated to radioresistance in GBMs, and in this case its activity was shown to be regulated post-transcriptionally by monitoring the phosphorylated state of the RELA protein^48^, providing an additional source of support for a second of the transcription factors identified here associated with clinical outcomes. In addition, we also identified RAR, which to our knowledge has not been linked to survival in GBM.

**Fig. 7.**
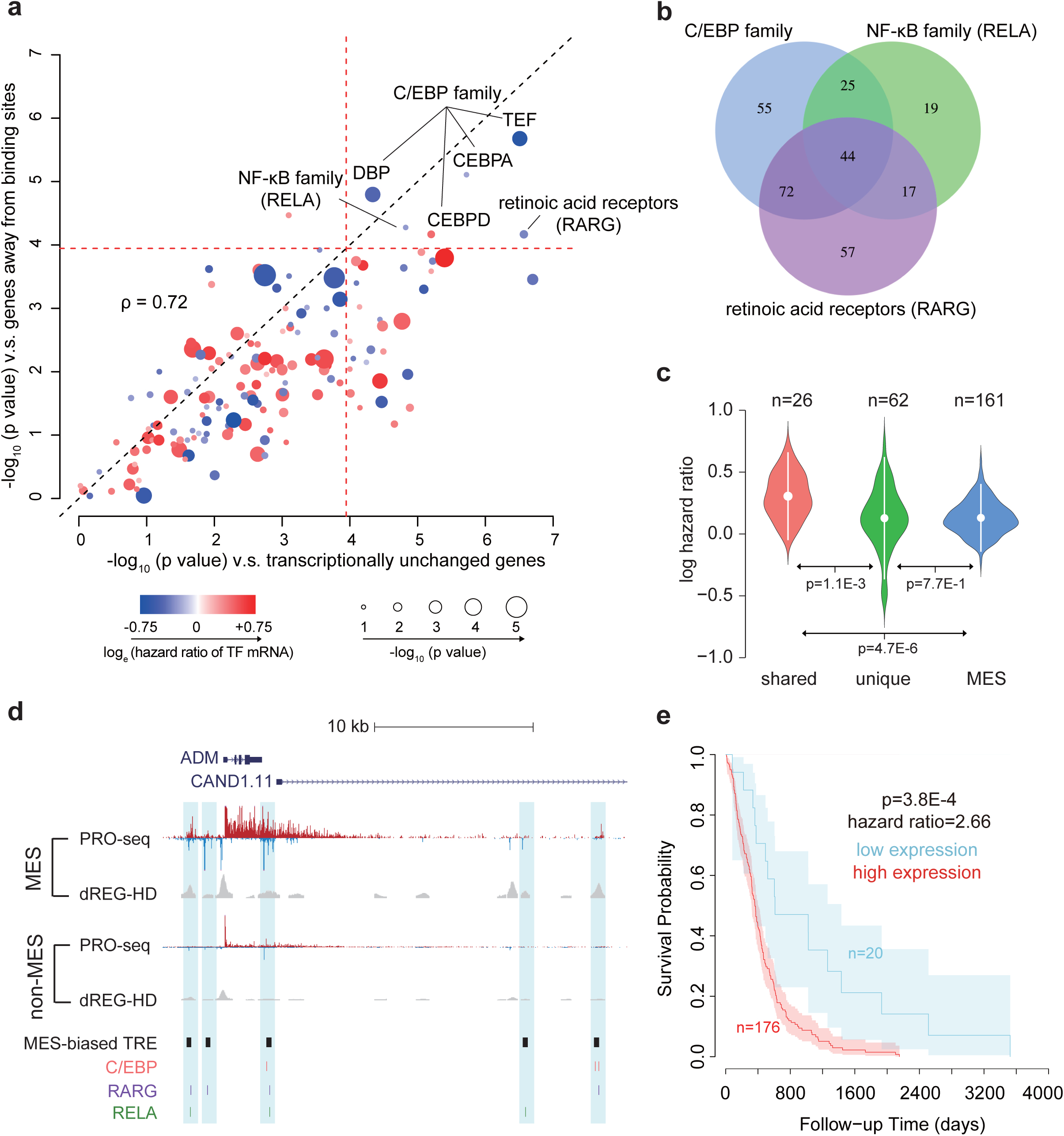
Transcription factors control survival associated pathways in GBM. **(a)** Scatter plots show the -log10 two-sided Wilcoxon rank sum test p value comparing the distribution of hazards ratios of target genes for each transcription factor and two groups of non-target control genes. One control represents genes that are close to transcription factor binding sites but do not change transcription levels in mesenchymal tumors (x-axis); the second control group represents mesenchymal up-regulated genes located distally (> 0.5 MB) from transcription factor binding sites (y-axis). The -log10 (p value) of association between transcription factor mRNA levels and survival is represented by the radius of the circle, and the natural log of the hazard ratio at higher mRNA levels is represented by the color. The dotted red line represents the Bonferroni adjusted a value. **(b)** Venn diagram shows overlap between the target genes of the three indicated survival associated transcription factors. **(c)** Violin plot shows the natural log of hazard ratios for target genes shared among (left) and unique to (center) the three transcription factors in **(b)**, and foi mesenchymal marker genes (right). Mean hazard ratios are shown by white dots and standard deviations are shown by bars. **(d)** Browser track of the ADM gene shows the average of RPM normalized (le)ChRO-seq signals and dREG-HD scores in mesenchymal and non-mesenchymal GBMs. Mesenchymal (MES)-biased TREs are highlighted in blue. The positions of MES-biased TRE and motifs of survival-associated transcription factors are shown on the bottom. **(e)** Kaplan–Meier plot shows the difference in overall survival between patients with high and low average expression level of shared target genes. The cutoff was determined based on the minimum p value in the difference between survival time using a Chi-squared test. Shaded regions mark the 95% confidence interval.

Surprisingly all three survival associated transcription factors regulated overlapping sets of putative target genes. Of four different combinations in which multiple transcription factors could regulate overlapping targets, three were more common than expected (*p* < 0.01; super exact test^53^; **Fig. 7b**; **Supplementary Fig. 28**), including 44 target genes that were shared among all three transcription factors. Target genes shared among all three transcription factors had significantly higher hazard ratios than unique target genes (**Fig. 7c,d**, *p* = 1.1e-3, Wilcoxon). Of the 26 shared targets for which hazards ratios were available, all were negatively correlated with survival, and eight were significantly associated with clinical outcomes on their own (a significant enrichment [*p* = 6e-4, Fisher’s exact test]), including *CCL20* (**Supplementary Fig. 29a**) and *ADM* (**Fig. 7d**), (*p* < 0.05, Chi-squared test) (**Supplementary Table 5**). High expression of both genes was associated with high risk regardless of subtype assignment, indicating that survival association of these transcription factors was not simply driven by enrichment in the mesenchymal subtype (**Supplementary Fig. 29b-c**). Moreover, differences in survival among these genes were not driven by IDH1 status (**Supplementary Fig. 30**). Gene ontology analysis found that targets of all three transcription factors were enriched for immune system process and stress responses (*p* < 1e-5, false discovery rate (FDR) corrected Fisher’s exact test, **Supplementary Table 6**). Taken together, our analysis suggests that C/EBP, RARG, and NF-κB work in concert to activate a shared regulatory program that controls inflammatory processes and correlates with poor clinical outcomes in GBM.

## Discussion

Nascent transcription is a promising approach for studying the molecular basis of complex disease because unstable RNAs provide deep insights into multiple stages of gene regulation. ChRO-seq allows maps of nascent transcription to be constructed in virtually any sample that maintains the integrity of protein-DNA interactions – even those in which RNA is highly degraded. ChRO-seq has important applications throughout the biomedical sciences in analyzing regulatory programs that contribute to solid tumors and other tissues which have proven challenging to study using existing molecular tools.

Our analysis of 20 primary tumors revealed several insights into transcriptional regulatory programs in malignant tissue. First, we report that enhancers in malignant tissue were surprisingly similar to DHSs in the tissue of origin. This finding suggests that regulatory programs in GBM often work within the confines of chromatin architecture that is established in the initiating cell. Regulatory programs were also similar to normal brain in PDXs, demonstrating that tumor initiating cells are able to reconstitute a diverse cell environment that bares surprising similarity to primary brain tissue. Yet how are malignant cell behaviors specified by cancer cells despite this similarity? We found a rare population of ectopic enhancers that resembled fetal tissues isolated from the nervous system, immune cells, and differentiated tumor cells. Our observations are consistent with models of tumorigenesis in which tumor cells reactivate regulatory programs that were similar in some respects to an earlier developmental stage^54^. These regulatory signatures derived from rare ectopic enhancers may have important prognostic value that can be exploited in future studies.

Our study highlights how transcription factors are responsible for coordinated changes in the expression of groups of genes that contribute to expression heterogeneity among tumors. ChRO-seq, like other run on technologies^55^, provides substantial information about the regulatory activities of transcription factors on chromatin that is independent of transcription factor expression levels. In support of our general approach, transcription factor candidates activating TREs in the stem-like regulatory program were similar to those reported previously to be sufficient for initiating tumors in a murine host^44^. Additionally, we used ChRO-seq data to identify transcription factors that establish differences in gene expression characteristic of reported GBM subtypes.

We report three transcription factors, C/EBP, RAR, and NF-κB, whose target genes were systematically correlated with poor clinical outcomes. Our work adds new transcription factors to the current literature, as well as additional support for the role of C/EBP in driving mesenchymal transformation^49^. NF-κB was previously associated with resistance to radiotherapy and involvement in mesenchymal transformation in GBMs^48^. Our present work builds on these studies to show that NF-κB activation has an unambiguous influence on clinical outcomes. Additionally, we found evidence that a third transcription factor, RAR, drives regulatory programs that contribute to survival in GBMs. Notably, post-transcriptional mechanisms are responsible for activating two of these three transcription factors, NF-κB and RAR. Thus insights reported here were possible only because ChRO-seq is a more direct indicator of transcription factor activity than other tools previously applied in GBM. As the pharmacology for targeting diverse transcription factor families develops, the transcription factors reported here, as well as our strategies for finding them, will become more useful in nominating targeted therapies.

## Online Methods

### Cell culture

Jurkat cells were grown in RPMI-1640 supplemented with 10% fetal bovine serum, 1X Penicillin/Streptomycin Antibiotic, 0.125 mg/ml Gentamicin Antibiotic at 37°C, 5% CO_2_. 1×10^6^ cells were centrifuged at 700 × g 4°C 5 min. The media was removed and the cells were rinsed with 1X PBS, centrifuged, and PBS was removed.

### Tissue collection and preparation

Glioblastoma-derived cells were prepared from freshly biopsied human tumors obtained with patient consent and approval by the Institutional Review Board at SUNY Upstate Hospital, Syracuse, NY. To establish patient-derived xenografts, small pieces of freshly resected gliomas were implanted subcutaneously in the flank of athymic nude (nu/nu) mice (Harlan Laboratories / Envigo, Indianapolis,IN) and serially passaged (mouse-to-mouse) 3 times for PDX-UMU88–02, 7 times for PDX-UMU89–08, and 57 times for PDX-88–04 p57, as previously described ^56,57^. To prepare chromatin pellets tissue samples were pulverized in a cell crusher. The Cellcrusher was chilled in liquid nitrogen. Frozen glioblastoma tissue (~ 100 mg) was placed in the Cellcrusher, the pestle is placed into the Cellcrusher, and the pestle was stuck with the mallet until the tissue was fractured into a fine powder.

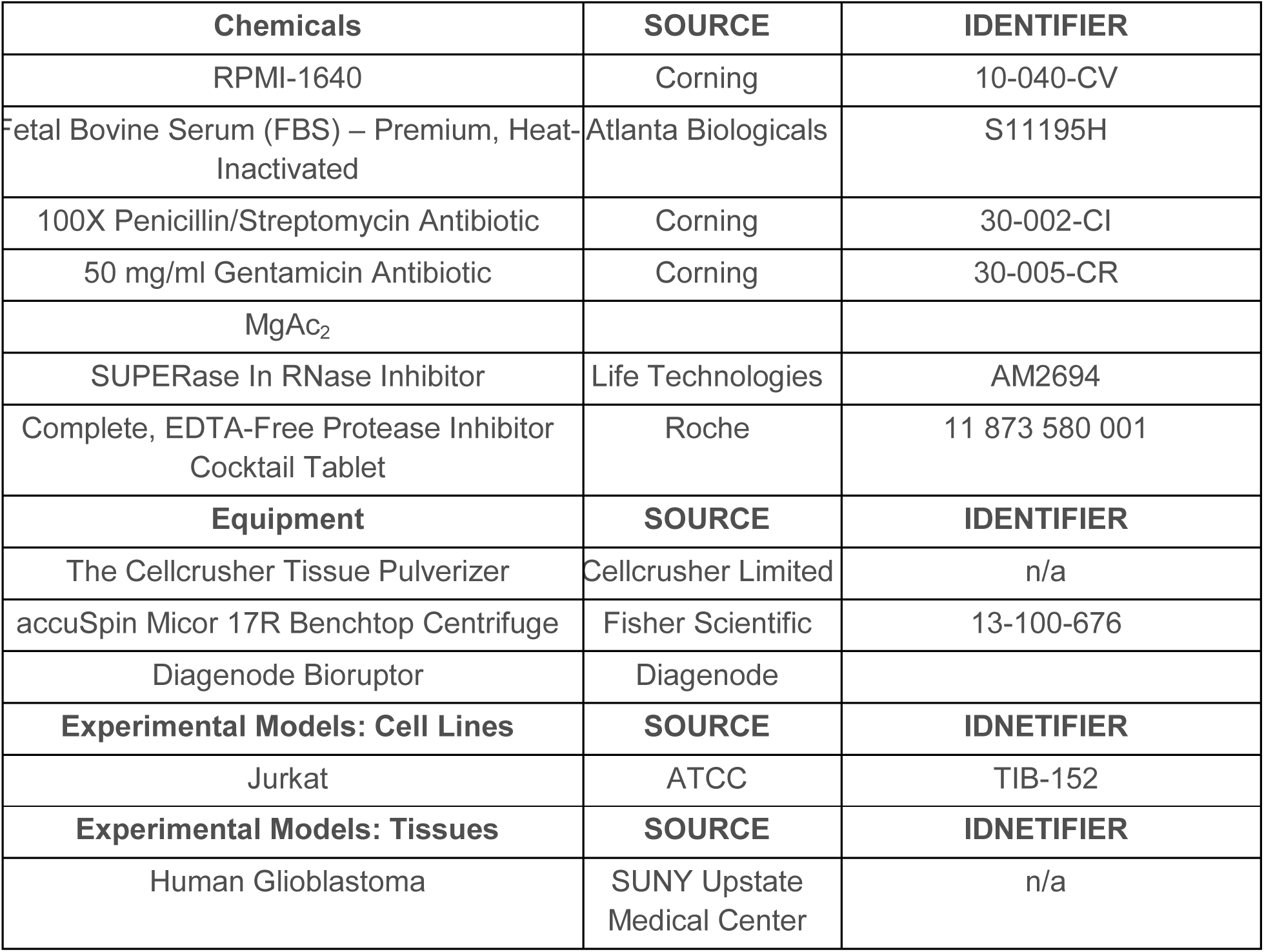
Table of key reagents in chromatin isolation

**Table of key reagents in chromatin isolation**
| Chemicals | SOURCE | IDENTIFIER |
| --- | --- | --- |
| RPMI-1640 | Corning | 10-040-CV |
| Fetal Bovine Serum (FBS) – Premium, Heat-Inactivated | Atlanta Biologicals | S11195H |
| 100X Penicillin/Streptomycin Antibiotic | Corning | 30-002-CI |
| 50 mg/ml Gentamicin Antibiotic | Corning | 30-005-CR |
| MgAc<sub>2</sub> |  |  |
| SUPERase In RNase Inhibitor | Life Technologies | AM2694 |
| Complete, EDTA-Free Protease Inhibitor Cocktail Tablet | Roche | 11 873 580 001 |
| Equipment | SOURCE | IDENTIFIER |
| The Cellcrusher Tissue Pulverizer | Cellcrusher Limited | n/a |
| accuSpin Micor 17R Benchtop Centrifuge | Fisher Scientific | 13-100-676 |
| Diagenode Bioruptor | Diagenode |  |
| Experimental Models: Cell Lines | SOURCE | IDNETIFIER |
| Jurkat | ATCC | TIB-152 |

| Experimental Models: Tissues | SOURCE | IDNETIFIER |
| --- | --- | --- |
| Human Glioblastoma | SUNY Upstate Medical Center | n/a |

### Chromatin isolation

The chromatin isolation was based on work first described in ref^27^. For chromatin (ChRO) isolation from cultured cells or tissue we added 1 ml of 1x NUN Buffer (0.3 M NaCl, 1M Urea, 1% NP-40, 20 mM HEPES, pH 7.5, 7.5 mM MgCl2, 0.2 mM EDTA, 1 mM DTT, 20 units/ml RNase Inhibitor (Life Technologies # AM2694), 1X Protease Inhibitor Cocktail (Roche # 11 873 580 001)). Samples were vigorously vortexed for one minute. An additional 500 μl of appropriate NUN Buffer was added to each sample and vigorously vortexed for an additional 30 seconds. For length extension chromatin (leChRO) isolation from cultured cells or tissue we added 1 ml of 1x NUN Buffer, as described previously, spiked with 50 units/ml RNase Cocktail Enzyme Mix (Ambion # 2286) in place of the RNase Inhibitor. The samples were incubated on ice for 30 minutes with a brief vortex every 10 minutes. Samples were centrifuged at 12,500 × g at 4°C for 30 minutes after which the NUN Buffer was removed from the chromatin pellet. The chromatin pellet was washed with 1 ml 50 mM Tris-HCl, pH 7.5 supplemented with 40 units/ml RNase Inhibitor (Life Technologies # AM2694), centrifuged at 10,000 × g, 4°C, for 5 minutes, and buffer discarded. The chromatin was washed two additional times. After washing, 100 μl of chromatin storage buffer (50mM Tris-HCl, pH 8.0, 25% Glycerol, 5mM MgAc2, 0.1mM EDTA, 5mM DTT, 40 units/ml RNase Inhibitor) was added to each sample. The samples were loaded into the Bioruptor and sonicated using the following conditions: power setting on high, cycle time of ten minutes with cycle durations of 30 seconds on and 30 seconds off. The sonication was repeated up to 3 times as needed to get the chromatin pellet into suspension. Samples were stored at - 80°C.

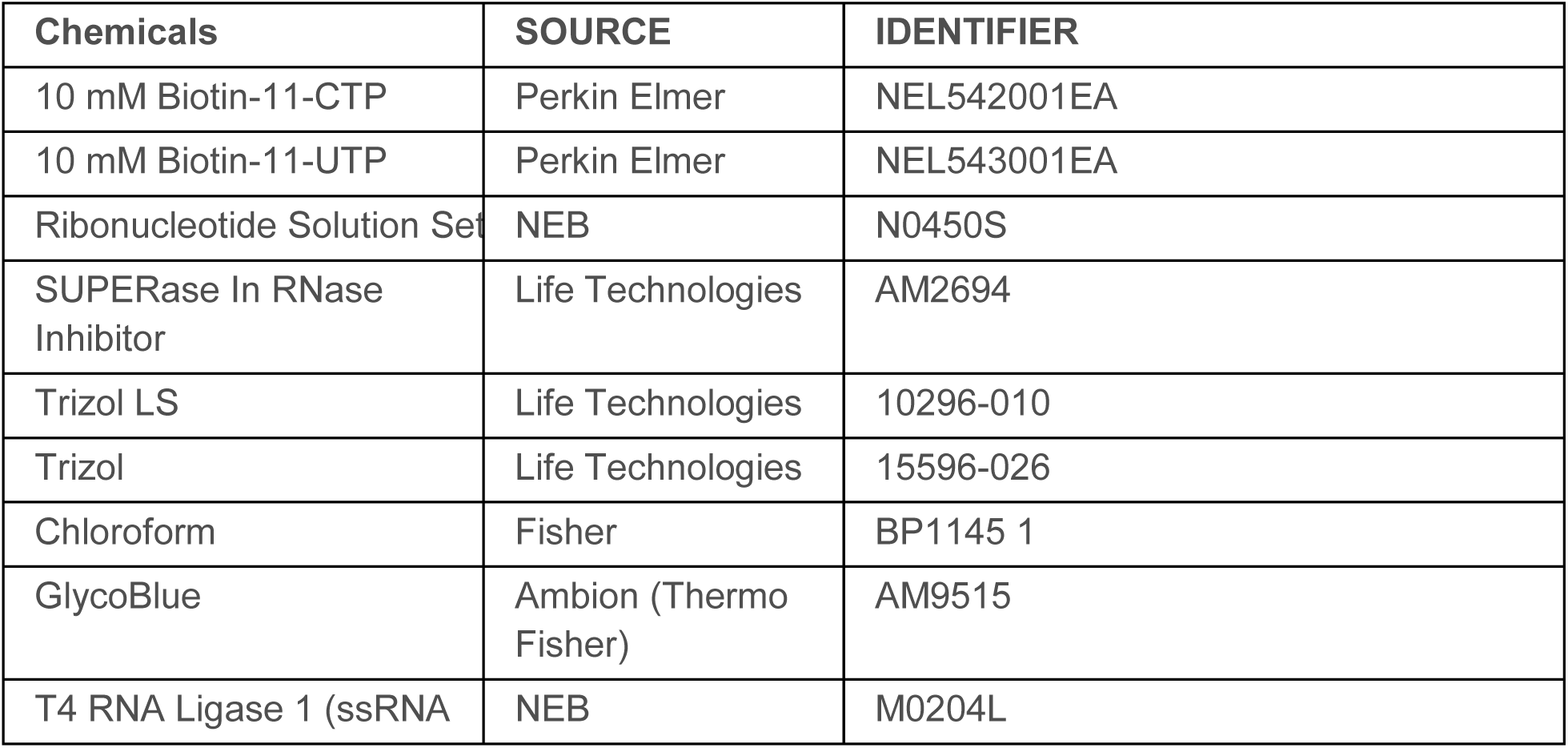
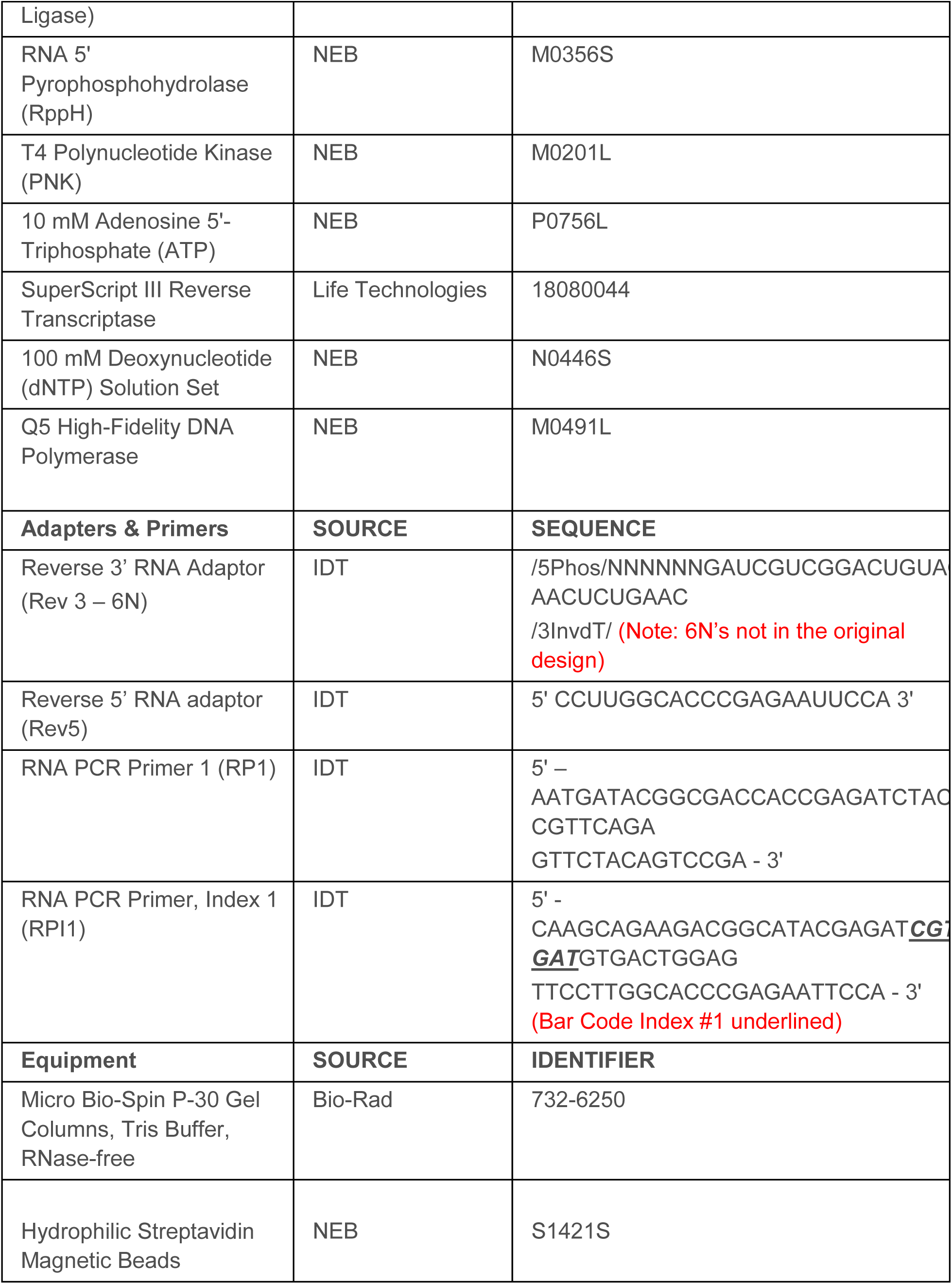
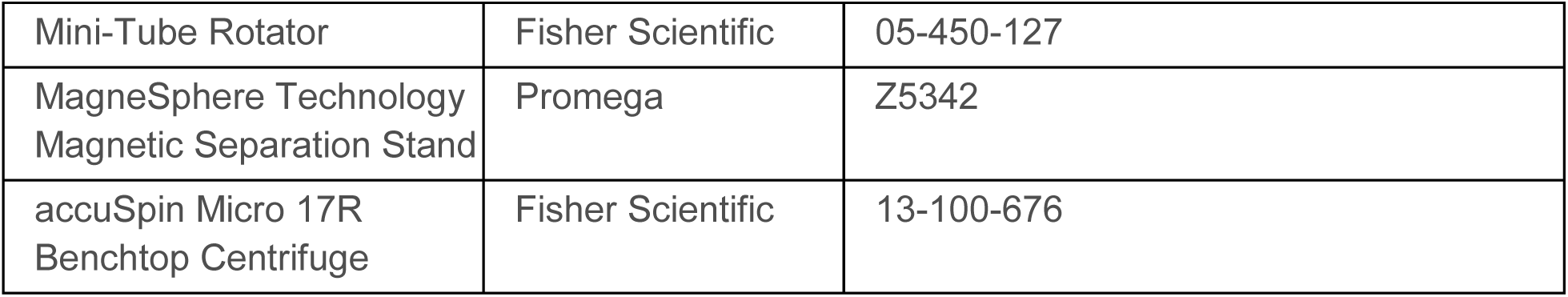
Table of Key Reagents in ChRO-seq

### Chromatin Run-On and sequencing (ChRO-seq) library preparation

After chromatin isolation, the chromatin run-on and sequencing library prep closely followed the methods described previously^28^. Briefly chromatin from 1×10^6^ Jurkat T-cells or 10–100 mg of primary glioblastoma or 100 mg of PDX in 100 chromatin storage buffer was mixed with 100 of 2x chromatin run-on buffer (10 mM Tris-HCl pH 8.0, 5 mM MgCl_2_,1 mM DTT, 300 mM KCl, 400 μM ATP (NEB # N0450S), 40 μM Biotin-11-CTP (Perkin Elmer # NEL542001EA), 400 μM GTP (NEB # N0450S), 40 μM Biotin-11-UTP (Perkin Elmer # NEL543001EA), 0.8 units/μl SUPERase In RNase Inhibitor (Life Technologies # AM2694), 1% Sarkosyl (Fisher Scientific # AC612075000)). The run-on reaction was incubated at 37°C for 5 minutes. The reaction was stopped by adding Trizol LS (Life Technologies # 10296–010) and pelleted with GlycoBlue (Ambion # AM9515) to visualize the RNA pellet. The RNA pellet was resuspended in DEPC treated water and heat denatured at 65°C for 40 seconds. In ChRO-seq, we digested RNA by base hydrolysis in 0.2N NaOH on ice for 8 minutes, which ideally yields RNA lengths ranging from 40 – 100 bases. This step was excluded from leChRO-seq. Nascent RNA was purified by binding streptavidin beads (NEB # S1421S) and washed as described^28^. RNA was removed from beads by Trizol and followed by the 3’ adapter ligation (NEB # M0204L). A second bead binding was performed followed by a 5’ de-capping with RppH (NEB # M0356S). The 5’ end was phosphorylated using PNK (NEB # M0201L) followed by a purification with Trizol (Life Technologies # 15596–026). A 5’ adapter was then ligated onto the RNA transcript. A third bead binding was then followed by a reverse transcription reaction to generate cDNA (Life Technologies # 18080–044). cDNA was then amplified (NEB # M0491L) to generate the ChRO-seq libraries which were prepared based on manufacturer's’ protocol (Illumina) and sequenced using Illumina NextSeq500 at the Cornell University Biotechnology Resource Center.

#### Mapping of ChRO-seq and leChRO-seq sequencing reads

We used our publicly available pipeline to align ChRO-seq and leChRO-seq data (https://github.com/Danko-Lab/utils/tree/master/proseq). Some libraries were prepared using adapters which contained a molecule-specific unique identifier (first 6 bp sequenced; denoted in Table 2), and for these we removed PCR duplicates using PRINSEQ lite ^58^. Adapters were trimmed from the 3’ end of remaining reads using cutadapt with a 10% error rate ^59^. Reads were mapped with BWA ^60^ to the human reference genome (hg19) plus a single copy of the Pol I ribosomal RNA transcription unit (GenBank ID# U13369.1). The location of the RNA polymerase active site was represented by a single base which denotes the 3’ end (ChRO-seq) or 5’ end (leChRO-seq) of the nascent RNA, which corresponds to the position on the 5’ or 3’ end of each sequenced read respectively. Mapped reads converted to bigWig format using BedTools ^61^ and the bedGraphToBigWig program in the Kent Source software package ^62^. Downstream data analysis was performed using the bigWig software package, available from: https://github.com/andrelmartins/bigWig. All data processing and visualization was done in the R statistical environment ^63^

#### Gene transcription activity quantification for ChRO-seq and leChRO-seq

We quantified transcription activity of ChRO-seq and leChRO-seq data using gene annotations from GENCODE v25 lift 37, expect for the cross-comparison with TCGA RNA-seq data, where we used GENCODE v22 lift 37 to match the annotation of TCGA data. We counted reads in the interval between 500 bp downstream of the annotated transcription start site to the end of the gene for comparisons. This window was selected to avoid counting reads in the pause peak near the transcription start site. We limited analyses to gene annotations longer than 1,000 bp in length.

### Molecular subtype classification

Transcriptional activity of characteristic genes for each GBM subtype (n = *23)* were quantified by the methods described above. Reads count from each sample are normalized by reads per million total reads count, followed by log2 transformation of pseudo count (RPM normalized reads count+1). The transformed read count is then centered with mean zero for each gene. The similarity between each sample was measured by Spearman’s rank correlation, and clustered using single link clustering. The subtype score was calculated by Pearson correlation with the centroid of corresponding subtype reported by^26^ (*n = 23*).

### Differential expression analysis (DESeq2) for annotated genes

Transcription activity of genes in each primary GBM / non-malignant brain were quantified by the methods described above. Patients clustered in each dominant subtype were treated as biological replicates (**Fig. 2b** and **Supplementary Table 3**). Two technical replicates of non-malignant brain were used as control. Differential expression analysis was conducted using deSeq2 (Love et al., 2014) and differentially expressed genes were defined as those with a false discovery rate (FDR) less than 0.05.

#### dREG-HD

*Overview*. We trained an epsilon-support vector regression (SVR) model that maps PRO-seq, GRO-seq, or ChRO-seq data to smoothed DNase-I-seq intensity values. Because dREG reliably identifies the location of transcribed TREs that are enriched for DHSs^19^, with its primary limitation being poor resolution, we limited the training and validation set to dREG sites. The SVR was trained to impute DNase-I values of the positions of interest based on its input PRO-seq data. The trained SVR can then be used to predict DNase-I-seq signal intensities in any cell type for which PRO-seq data is available. To identify the location of transcribed DNase-I hypersensitive sites (DHSs) we developed a heuristic method to identify local maxima in imputed DNase I-seq data. A detailed description of these tools is provided in the following sections. The source code for the R package of dREG-HD is available from https://github.com/Danko-Lab/dREG.HD.git.

*Training the dREG-HD support vector regression model*. PRO-seq data was normalized by the number of mapped reads and was summarized as a feature vector consisting of ±1800 bp surrounding each site of interest. In total, 113,568 sites were selected, and were divided into 80% for training and 20% for validation. Parameters for the feature vector (e.g., window size) were selected by maximizing the Pearson correlation coefficients between the imputed and experimental DNase-I score over the holdout validation set used during model training (**Supplementary table 4**). We fit an epsilon-support vector regression model using the Rgtsvm R package^64^.

We optimized several tuning parameters of the model during training. We tested various kernels, including linear, Gaussian, and sigmoidal. Only the Gaussian kernel was able to accurately impute the DNase-I profile. Experiments optimizing the window size and number of windows revealed that feature vectors with the same total length but different step size result in similar performance on the validation set, but certain combinations with fewer windows achieved much less run time in practice. The feature vector we selected for dREG-HD used non-overlapping windows of 60bp in size and 30 windows upstream and downstream of each site, and resulted in near-maximal accuracy and short run times on real data. To make imputation less sensitive to outliers, we scaled the normalized PRO-seq feature vector during imputation such that its maximum value is within the 90th percentile of the training examples. This adjustment makes the imputation less sensitive to outliers and improves the correlation and FDR by 4% and 2%, respectively.

The optimized model achieved a log scale Pearson correlation with experimental DNase-I seq data integrated over 80bp non-overlapping windows within dREG regions of 0.66 at sites held out from the K562 dataset on which dREG-HD was trained and 0.60 in a GM12878 GRO-seq dataset that was completely held out during model training and parameter optimization (**Supplementary Fig. 9**).

*Curve fitting and peak calling*. The imputed DNase-I values were subjected to smoothing and peak calling within each contiguous dREG region. To avoid effects on the edge of dREG regions, we extended dREG sites by ±200bp on each side before peak calling. We fit the imputed DNase-I signal using smoothing cubic spline. We defined a parameter, the knots ratio, to control the degree to which curve fitting smoothed the dREG-HD signal. The degree of freedom (λ) of curve fitting for each extended dREG region was controlled by knots ratio using the following formula.

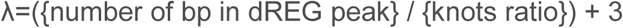

This formulation allowed the equivalent degrees of freedom to increase proportionally to the length of the dREG peak size, but kept the value of the knots ratio higher than a cubic polynomial.

Next we identified peaks in the imputed dREG-HD signal, defined as local maxima in the smoothed imputed DNase-I-seq profiles. We identified peaks using a set of heuristics. First, we identify local maxima in the dREG-HD signal by regions with a first order derivative of 0. The peak is defined to span the entire region with a negative second order derivative. Because dREG-HD is assumed to fit the shape of a Guassian, this approach constrains peaks to occur in the region between ±σ for a Gaussian-shaped imputed DNase-I profile. We optimized curve fitting and peak calling over two parameters: 1) knots ratio and 2) threshold on the absolute height of a peak. Values of the two parameters were optimized over a grid to achieve a balance between sensitivity and false discovery rate (FDR). We chose two separate parameter combinations: one ‘relaxed’ set of peaks (knots ratio=397.4, and background threshold=0.02) that optimizes for high sensitivity (sensitivity=0.94 @ 0.17 FDR), and one stringent condition (knots ratio=1350 and background threshold=0.026) that optimizes for low FDR (sensitivity=0.79 @ 0.07FDR).

*Validation metric and genome wide performance*. We used genomic data in GM12878 and K562 cell lines to train and evaluate the performance of dREG-HD genome-wide. Specificity was defined as the fraction of dREG-HD peaks calls that intersect with at least one of the following sources of genomic data: Duke DNase-I peaks, UW DNase-I peaks, or GRO-cap HMM peaks. Sensitivity was defined as the fraction of true positives, or sites supported by all three sources of data that also overlapped with dREG. To avoid creating small peaks by an intersection operation, all data was merged by first taking a union operation and then by finding sites that are covered by all three data sources. All dREG-HD model training was performed on K562 data. Data from GM12878 was used as a complete holdout dataset that was not used during model training or parameter optimization.

*Metaplots for dREG and dREG-HD*. Metaplots were generated using the bigWig package for R with the default settings. This package used a subsampling approach to find the profile near a typical site, similar to ref^65^. Our approach samples 10% of the peaks without replacement. We take the center of each dREG-HD site and sum up reads by windows of size 20bp for total of 2000 bp in each direction. The sampling procedure is repeated 1000 times, and for each window the 25% quartile (bottom of gray interval), median (solid line), and 75% quartile (top of tray interval) were calculated and displayed on the plot. Data from all plots were generated by the ENCODE project ^42^.

#### Data processing for calling DNase-I hypersensitive sites and dREG-HD sites

We reprocessed all DNase-I-seq data and identified DNase-I hypersensitive sites (DHSs) using a uniform pipeline. We retrieved mapped reads from either ENCODE or Epigenome roadmap projects aligned to hg19. We called peaks in individual biological replicates, 921 samples in total, using MACS2 ^66^ and Hotspot. To group DHSs for each cell and tissue type with high confidence, we took the union of peaks (bedtools merge) from biological replicates followed by intersecting peaks called by Hotspot and MACS2. Lastly since peaks resulted from intersection may be too narrow and hence become missed during downstream intersection operations, we expanded all short peaks (<150bp) to 150bp from the peak center. Analyses involving individual replicates, in **Supplementary Fig.11**, use only peaks called by MACS2.

ChRO/leChRO-seq data was mapped to hg19 as described above. dREG score was thresholded at 0.7 to generate dREG peak regions for dREG-HD run. dREG-HD runs were done at the stringent condition, except for analysis of subtype biased TREs, where we used dREG-HD sites called at relaxed condition.

#### Mutual information analysis

We used mutual information to compare the similarity between TREs observed in any pair of DHS or dREG-HD datasets. DHSs or dREG-HD peaks of sample involved in the comparison were merged in order to construct a sample space in which two or more samples would be compared. Each dataset was then summarized as a random variable, represented by a zero-one vector in which each element represents a TREs in the sample space, and takes a value of 1 if it intersects with that peak and 0 otherwise. We calculated the mutual information between two random variables, X and Y, using the formula below:

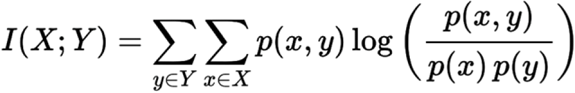

#### Comparison between tumor and reference brain tissues and cell lines

We selected brain-related samples from uniformly processed DHSs and categorized the reference dataset by sample origin, namely normal adult brain tissues (globus pallidus, midbrain, frontal cortex, middle frontal gyrus, cerebellum and cerebellar cortex), primary brain cells (astrocyte of the hippocampus, astrocyte of the cerebellum, and normal human astrocytes), and GBM cell lines (A172, H54 and M059J).

#### Mutual information heatmap and clustering analysis

To compare the similarity between the dREG-HD sites in each query samples and DHSs in each reference samples (**Fig. 3c**), we computed the pairwise mutual information between each pair of dREG-HD and DHSs (as described above) on the sample space defined by merged peaks among all samples included in the analysis. We noted a systematic bias in the distribution of mutual information across query samples that appeared to reflect data quality and sequencing depth in either ChRO-seq or DNase-I-seq data. To correct this bias, we normalized the mutual information of each query sample with respect to the sum of mutual information for that query sample.

Among multiple samples normalizing the mutual information metric is more complicated. We devised an approach that was used in **Supplementary Fig. 14**. We consider a square matrix with rows and columns representing each sample. Each element in this matrix represents the mutual information between a pair of samples. Our objective is to center the mutual information across each row or column while preserving the rank order and range of mutual information. We accomplished this using the following algorithm, which is similar to ^67^, but guarantees symmetry:

~~~
#matrix centering algorithm
WHILE convergence criterion does not meet
FOR i from 1 to number of columns
            current mean<-mean of ith column
            ith row <-ith row – current mean
            ith column <-ith column – current mean
END FOR
END WHILE
~~~

The convergence criterion was defined as the maximum of the absolute value of element-wise difference between matrix returned from previous two consecutive runs. Although there is no mathematical guarantee of convergence, this approach converged typically after four cycles with the datasets that we used. After centering the matrix was clustered using the ward.D2 clustering algorithm implemented in the heatmap function in R.

#### TRE clustering analysis

We analyzed the activation pattern across TREs, using the same definition of sample space described in the mutual information analysis (above). We assigned two states to each TRE, active if intersected dREG-HD/ DHS, and inactive if otherwise. The Jaccard distance was used to quantify the similarity between two samples or between two potential TREs. Clustering across samples (columns) and across TREs (rows) was done using ward.D2 method. To reduce the influence of noise on the clusters, we limited analysis to TREs that were activated in at least two query samples but less than 6 brain-related reference samples (16 samples in total).

#### taTRE enrichment test and clustering into regulatory programs

taTREs were defined as TREs from primary GBM / PDX that do not intersect with any dREG-HD peaks from our non-malignant brain control nor with DHSs found in normal brain tissues (including globus pallidus, midbrain, frontal cortex, middle frontal gyrus, cerebellum and cerebellar cortex). These taTREs represent a stringent subset enriched for TREs associated with the malignant phenotypes observed in brain tumors. dREG-HD sites or DHSs that overlapped with ENCODE consensus hg19 blacklist regions were excluded from analysis.

The majority of taTREs intersected DHSs in one or more reference ENCODE and Epigenome Roadmap samples (**Fig.3a**). We devised a statistical test to determine whether the observed number of intersections with each reference sample is significantly higher than expected by chance. We generated a null distribution by sampling DHSs with replacement from all TREs found in reference samples, controlling for the distribution of uniqueness (i.e., the number reference samples which each taTRE intersects) of taTREs from a particular GBM / PDX. The simulation was run for 10^5^ times for each sample, each simulation drawing the same number of taTREs observed in that sample. We selected tissues with a stringent statistical significance cutoff of p(X_null_ > x_observed_) ≤ 1/10^4^. Reference samples that showed significant enrichment in at least one third of (≥8) GBM or PDX were chosen as taTRE-associated references for downstream analysis.

In total 50 significant taTRE-enriched reference samples were clustered by methods described in the *TRE clustering analysis* section. Fold of enrichment was calculated as the x_observed_ / E[X_null_]. The dendrogram was cut down to three clusters. DHS regions that show up in more than half of reference samples in each cluster were picked as representative DHS driving a regulatory program that is characteristic for that cluster. taTREs overlapping these representative DHSs unique to each cluster were selected for downstream analysis.

#### Motif enrichment analysis of tumor-associated TREs

*Defining genomic regions for motif enrichment comparison*. taTREs from the group indicated in the **Supplementary Fig. 15** (positive set) were compared against normal brain TRE (background set). Normal brain TREs (nbTRE) were constructed from the dREG-HD sites that intersect with active DHSs peaks in the adult normal brain. For the positive and background sets we selected the center of peaks and then extended by 150bp from the center. We subsampled background peaks to construct >2,500 GC-content matched TREs before scanning for motif enrichment.

*Motif enrichment analysis*. We used the R package rtfbsdb to search for motifs that show enrichment in each primary GBM^47^. We focused on 1,882 human transcription factor binding motifs from the CisBP database^68^. When scanning genomic regions of interest, we used TFBSs having a log_e_-odds score ≥7 in positive and background sets, with scores obtained by comparing each representative motif model to a second-order Markov background model. Motif enrichment was tested using Fisher’s exact test. To account for potential bias resulted from difference in GC-content between positive and background sets, we ran statistical test on 50 independently subsampled GC-matched dREG-HD regions, and summarized the p values and the fold enrichment across background sets by the median across samples. To search for motif enrichment across 1,882 human transcription factor binding motifs in each patient (all taTRE against all normal brain TRE), we define criterion as follows: 1) The fold of enrichment was greater than 1, 2) the enrichment was robustly significant to changes in the GC matched background set (median *p* < 0.05/1882), 3) the positive sets have at least 10 sites with log_e_-odds score ≥7, 4) the transcription factor was transcribed with at least 2 ChRO/leChRO-seq reads in its gene body.

*Summarizing motif enrichment statistics across patients*. Motifs that were enriched in at least one primary GBM (all taTRE against all normal brain TRE) were chosen for downstream analysis. The enrichment statistics of three regulatory modules-taTREs were also summarized by median over the patients that show significant enrichment for the motif. Lastly, for each transcription factors with multiple motif IDs, we reported the one with the most significantly enrichment in all taTREs over nbTREs.

#### Motif enrichment analysis of subtype-biased TREs

*Defining subtype-biased TREs*. To search for TREs that differentially activated or repressed in each subtype, we rely on measuring the change of the nascent RNA in the TRE regions. We merged dREG-HD sites called using the relaxed setting across 23 samples. We summed up the reads count of leChRO/ChRO-seq of each merged dREG-HD sites extended by 250bp from the center. TREs in patients of the subtype of interest (**Supplementary Table 4**) were compared against those of the rest three subtypes. Differential expression analysis was conducted using DESeq2^36^, and *subtype-biased TREs* are defined as those differentially transcribed with a false discovery rate (FDR) less than 0.01.

*Defining genomic regions for motif enrichment comparison*. Up or down-regulated subtype-specific TREs (positive set) were compared against TREs that did not show significant differential transcription (FDR DESeq2 *p* > 0.1) (background set). We scanned the dREG-HD regions extended by 150bp from the center of TREs, and subsampled background peaks to construct >2,500 GC-content matched TREs before scanning for motif enrichment.

*Motif enrichment analysis of subtype-biased TREs*. Motif enrichment analysis of *subtype-biased TREs* was done similarly to that for taTREs. The only minor difference was the strategy of filtering 1,882 human transcription factor binding motifs in each subtype. Criterion 1, 2, and 3 were identical to that for taTREs, while we modified the last criterion on transcription level of transcription factor to accommodate for replicates used for each subtype. For transcription factor motifs enriched in up-regulated subtype-biased TREs, we required at least 2 ChRO/leChRO-seq reads in its gene body in all samples of the subtype of interest. For those enriched in down-regulated subtype-biased TREs, we require at least 2 ChRO/leChRO-seq reads in its gene body in all samples of the rest three subtypes.

#### Motif clustering by genomic positions

Because we are not able to rigorously distinguish between paralogous transcription factors that share similar DNA binding specificities, we developed a method of clustering them based on their occurrence in the context of genomic regions. We first scanned motifs enriched over genomic regions defined by the positive set. In clustering motifs enriched in taTREs, we used the taTREs merged over 20 primary GBMs as the positive set; for motifs enriched in subtype biased TREs, we used the corresponding subtype biased TRE in which the motifs were enriched as the positive set. We defined the presence of TFBSs for loci (stand-specific) having a log_e_-odds score ≥7 in positive and background sets, and absence otherwise, with scores obtained by the method described in the section *Motif enrichment analysis of taTRE*. The Spearman’s rank order correlation coefficients were computed for each pair of transcription factors, based on their presence/absence pattern across TFBSs of all motifs of interest. Heatmaps were generated using agglomerative hierarchical clustering using the ward.D2 method.

#### Validation of regulation between transcription factors and target genes

*Associating transcription factors to target genes*. We associated transcription factors to target genes by first identifying its target TREs, and then search for target genes based on location of these TREs. To identify target TREs, we scanned "relaxed dREG-HD all GBM” regions, extended by 150bp from the center, using itself as the second-order Markov background model. For each subtype-specific transcription factor, we defined its binding sites as 1) ssTREs that undergo differentially transcription in the same subtype, and 2) have a log_e_-odds score ≥7 for at least one corresponding motif ids that also showed enrichment (p<0.05). This subset of TREs represents the potential binding and regulating sites of the TF of interest, referred to as query TREs. We use stringent heuristics link the query TREs to target genes in order to reduce false positive links. TREs were linked to putative target genes if: 1) the annotated transcriptional start site of the genes is the first two closest to the query TRE and within 50kb, and 2) the gene is differentially transcribed (FDR corrected DESeq2 *p* < 0.05) in the same direction as the query TRE.

*Defining the background set of non-target genes*. We defined background non-target genes of each transcription factor as those distal from (>0.5 Mb) the query TRE, but which show similar changes in transcription as that of target genes (to control for subtype). We required non-target genes had a transcription start site >0.5Mb from the closest query TRE. To match changes in transcription between target and non-target genes, we subsampled half of the genes away from query TREs and differentially transcribed (p<0.05) in the same direction as that of target genes without replacement, such that the distribution of log2 of fold change in transcription was insignificant (two-sided Wilcoxon *p* > 0.2).

*Validation of association between transcription factors and target genes*. To validate of our approach associating transcription factors to target genes, we compared the co-expression of target genes to that of background non-target genes. Specifically, we used the RPKM normalized TCGA RNA-seq data from 174 GBM patients downloaded from https://portal.gdc.cancer.gov/, and used the Spearman’s rank correlation to measure the degree of co-expression. To avoid the potential co-expression that might be artificially enriched in target genes due to higher chance of being located in adjacent positions of the genome, we masked the correlations coefficients between adjacent genes. We computed the significance for target genes to have higher co-expression using one-sided Wilcoxon rank-sum test.

*Quantifying the association between the transcription level of transcription factors and its target genes*. We used the RPKM normalized TCGA RNA-seq data from 174 GBM patients, and used the Spearman’s rank correlation to measure the monotonic relation between the transcription level of transcription factors and the putative target genes. We compared the difference between the distribution of correlation coefficients for target and non-target genes using the Wilcoxon rank-sum test and derive the two-sided *p* value.

#### Identification of transcription factors driving survival-associated programs

For each subtype-specific transcription factor, we identified the target genes as described above, and compared the hazard ratio of the target genes with that of non-target genes. We defined two sets of background based on non-target genes: 1) the closest genes whose transcription start site was also within 50 kb to the query TRE, but whose transcription unchanged across the samples representing that subtype (*p* > 0.2, **Fig. 7a**, x axis), and 2) genes differentially transcribed (*p* < 0.05) in the same direction as target genes, whose transcription start sites were 0.5Mb away from the closest query TRE (**Fig. 7a**, y axis). The clinical data, the scaled mRNA abundance level of 11,861 genes across 202 GBM patients, and unified over three microarray platforms, was downloaded from TCGA (https://tcga-data.nci.nih.gov/docs/publications/gbmexp/unifiedScaled.txt)^26^. We computed the hazard ratio of each gene by fitting a Cox proportional hazards regression model for survival time of patients with expression level in upper 25% of transcription levels over those with lower 25%. This ensures that all genes were fit for the regression model using the same balanced number of patients. We used the Wilcoxon test to compare the distribution of hazard ratios of target genes and background genes, and derived a two-sided *p* values for each background set.

The hazard ratio of analysis for individual transcription factors in **Fig. 7a** and **Supplementary Fig. 27a-c**, and target genes of survival-related transcription factors in **Fig. 7e**, **Supplementary Fig. 27d-f** and 30, were determined by the same regression model. The difference was that, instead of using the upper and lower quartiles as the cutoff, we reported the hazard ratio at the threshold between 0.1 quantile and 0.9 quantile that gave the largest difference between survival times. This difference was calculated by two-sided *p* value from Chi-squared test. This ensured that we reported the largest possible difference in survival time for each individual gene.

## Acknowledgements

We thank M. Viapiano and P. Sethupathy for valuable comments on the manuscript and other members of the Danko, Kwak, and Lis labs for valuable discussions. This research was supported, in part, by US National Institutes of Health (NIH) grants HG009309 to C.G.D. and H.K. and GM25232 to JTL and by the Walbridge Foundation for Brain Cancer Research. The content is solely the responsibility of the authors and does not necessarily represent the official views of the NIH or the Walbridge Foundation.

## Author Contributions

TC, ZW, and CGD analyzed the data. EJR, GB, and HK performed molecular experiments. HK conceived of a chromatin run-on, with input from LJC and JTL. HHS selected tumors for analysis from the GBM tissue bank. RJC and HHS completed the pathologic analysis. LSC and HHS dissected GBM-15–90 brain tissue. SLL runs the GBM tissue bank and performed the murine xenograft experiments. Data collection and analysis was supervised by CGD. The manuscript was written by CGD and TC, with input from the other authors.

## Author information

All ChRO-seq and leChRO-seq data is currently being deposited into the database of genotypes and phenotypes (dbGaP). Processed data will be deposited into Gene Expression Omnibus in hg19 and hg38 coordinate systems. All data analysis scripts and custom software will be distributed publicly on GitHub.

